# Membrane architecture and adherens junctions contribute to strong Notch pathway activation

**DOI:** 10.1101/2021.05.26.445755

**Authors:** Julia Falo-Sanjuan, Sarah J. Bray

## Abstract

The Notch pathway mediates cell-to-cell communication in a variety of tissues, developmental stages and organisms. Pathway activation relies on the interaction between transmembrane ligands and receptors on adjacent cells. As such, pathway activity could be influenced by the size, composition or dynamics of contacts between membranes. The initiation of Notch signalling in the *Drosophila* embryo occurs during cellularization, when lateral cell membranes and adherens junctions are first being deposited, allowing us to investigate the importance of membrane architecture and specific junctional domains for signaling. By measuring Notch dependent transcription in live embryos we established that it initiates while lateral membranes are growing and that signalling onset correlates with a specific phase in their formation. However, the length of the lateral membranes *per se* was not limiting. Rather, the adherens junctions, which assemble concurrently with membrane deposition, contributed to the high levels of signalling required for transcription, as indicated by the consequences from depleting α-Catenin. Together, these results demonstrate that the establishment of lateral membrane contacts can be limiting for Notch trans-activation and suggest that adherens junctions play an important role in modulating Notch activity.

## Introduction

The Notch pathway is a cell-cell signalling pathway conserved across animals with widespread roles in development, homeostasis and disease. Following interaction between the transmembrane Notch receptor(s) and ligands of the Delta or Serrate/Jagged families in adjacent cells, Notch is cleaved and the intracellular domain (NICD) translocates to the nucleus, where it regulates transcription of target genes. This limits signalling to cells that are directly in contact, although in some cases the contacts may occur though long cellular processes that extend between cells at a distance ***(Hunter et al. 2019; Boukhatmi et al. 2020)*.** Tissue geometry and the nature of the cell contacts will thus impact on the levels as well as the duration of signal that a cell receives from its neighbours ***(Shaya et al. 2017)*.** Elucidating the contributions from tissue architecture to Notch signalling will therefore be important to understand how signalling is effectively deployed in the different processes it controls.

An example where the acquisition of cell architecture may be important is during cellularization in *Drosophila*. Profound morphological changes take place at this stage, which corresponds to the onset of Notch signalling in the mesectoderm, a stripe of cells located between the mesoderm and mesectoderm that gives rise to the future midline of the ventral nerve cord ***(Nambu et al. 1990; Morel and Schweisguth 2000; Morel et al. 2003)*.** Prior to nuclear cycle 14 (nc14), the *Drosophila* embryo is a syncytium - the nuclei divide but are not separated by membranes. During nc14 membranes ingress to build intracellular membranes surrounding each nucleus, creating ~6000 cells, a process referred to as cellularization ***(Foe and Alberts 1983; Lecuit and Wieschaus 2000; Lecuit et al. 2002)*.**

In analyzing the real-time response of two well-characterized Notch responsive mesectodermal enhancers - *m5/m8* from *E(spl)-C* and the mesectodermal enhancer from *single-minded* (*sim*) ***(MartÍn-Bermudo et al. 1995; Cowden and Levine 2002; Zinzen et al. 2006; Hong et al. 2013)*** - during nc14, we observed that Notch dependent transcription was first detectable at 30 min into nc14 ***(Falo-Sanjuan et al. 2019)*.** This differs from other enhancers active at this stage, which exhibit high levels of activity from the beginning of nc14 ***(Garcia et al. 2013; Bothma et al. 2014; Bothma et al. 2015; Lim et al. 2017)*.** Ectopic production of NICD, which does not depend on membrane release and trafficking, was sufficient to produce earlier *m5/m8* and *sim* activity, suggesting that factors downstream of NICD production, such as co-activators or chromatin landscape, are not limiting at this stage. Based on the timing, and the fact that the two Notch responsive enhancers had similar onset times, we hypothesized that signalling was initiated at a defined morphological stage, releasing NICD to initiate transcription ***(Falo-Sanjuan et al. 2019)*.** This could arise due to the formation of lateral membrane and cell junctions or to other changes that occur during cellularization, such as alterations in nuclear size and shape ***(Brandt et al. 2006; Pilot et al. 2006)*.**

The timing and progression of cellularization is coordinated by two zygotically expressed proteins, Slam and Nullo, which are localized to the basal domain of the ingressing membranes ***(Hunter and Wieschaus 2000; Lecuit et al. 2002)*.** Slam activates Rho signaling by recruiting RhoGEF2 to the prospective basal domain, where it promotes actin polymerization and actomyosin contractility, resulting in furrow invagination ***(Wenzl et al. 2010)*.** Likewise, Nullo stabilizes the lateral furrows by regulating endocytic dynamics, which helps localize proteins to the basal junctions and impacts on actomyosin contractility ***(Sokac and Wieschaus 2008a; Sokac and Wieschaus 2008b)*.** As cellularization proceeds, cadherin-catenin complexes are assembled into first basal and then apical Adherens Junctions (AJs) that delimit the apical and basolateral domains ***(Hunter and Wieschaus 2000; Kramer 2000)*.** This step-wise progression of lateral membrane growth and junction formation offers a unique opportunity to explore the relationship between lateral membrane growth and competence for Notch signalling. We hypothesized that Notch signalling cannot initiate until the appropriate membrane domains are formed and matured, so that the ligand and receptor can be appropriately juxtaposed. However, it has also been proposed that autocrine signalling can occur, whereby productive interactions occur between the ligand and receptor in the same cell, either on the cell surface or on intracellular membrane vesicles (*e.g.* endosomes) ***(Coumailleau et al. 2009; Nandagopal et al. 2019)*.** By investigating the onset of signalling during cellularization we aim to resolve these models.

To distinguish the different models to explain signalling onset, we have assessed which processes during cellularization can affect the timing and/or levels of Notch dependent transcription in the mesectoderm. We find that Notch and Delta are present on the ingrowing lateral membranes and that signalling onset is highly correlated with membrane growth, but not with nuclear shape changes. Furthermore, the results suggest that the presence of lateral membranes *per se* is not sufficient for activation, and that high levels of signalling also require the establishment of cellular junctions, whose integrity regulates the turnover of Notch at the membrane. Whether the junctions contribute directly or indirectly to the signalling capabilities remains to be established, but the evidence clearly points to membrane morphogenesis, and the establishment of signalling competent membrane domains, as a key determinant for the initiation of Notch signalling in the embryo.

## Results

### Initiation of Notch-Delta signaling coincides with growth of lateral membranes

Notch dependent transcription in the mesectoderm is first detected approximately 30-35 minutes after the mitosis that marks the start of nc14, as illustrated by activity of *m5/m8* enhancer. This behaviour differs from that of other enhancers at this stage, which are active from the start of nc14 (**Fig.** 1**AB**). High levels of NICD can bypass the temporal restriction, directing expression much earlier ***(Falo-Sanjuan et al. 2019)*,** arguing that the Notch-regulated enhancers are competent to respond and that another step, besides enhancer accessibility, is limiting transcription onset. Nuclear maturation and cellularization are two developmental processes that occur during nc14 and could potentially govern the onset of Notch dependent transcription in a direct or indirect manner. We therefore began by characterizing how each of these processes related to the timing of transcription, measured using the MS2/MCP system ***(Garcia et al. 2013)*** to detect activity from the Notch-dependent *m5/m8* enhancer in real time.

**Figure 1.**
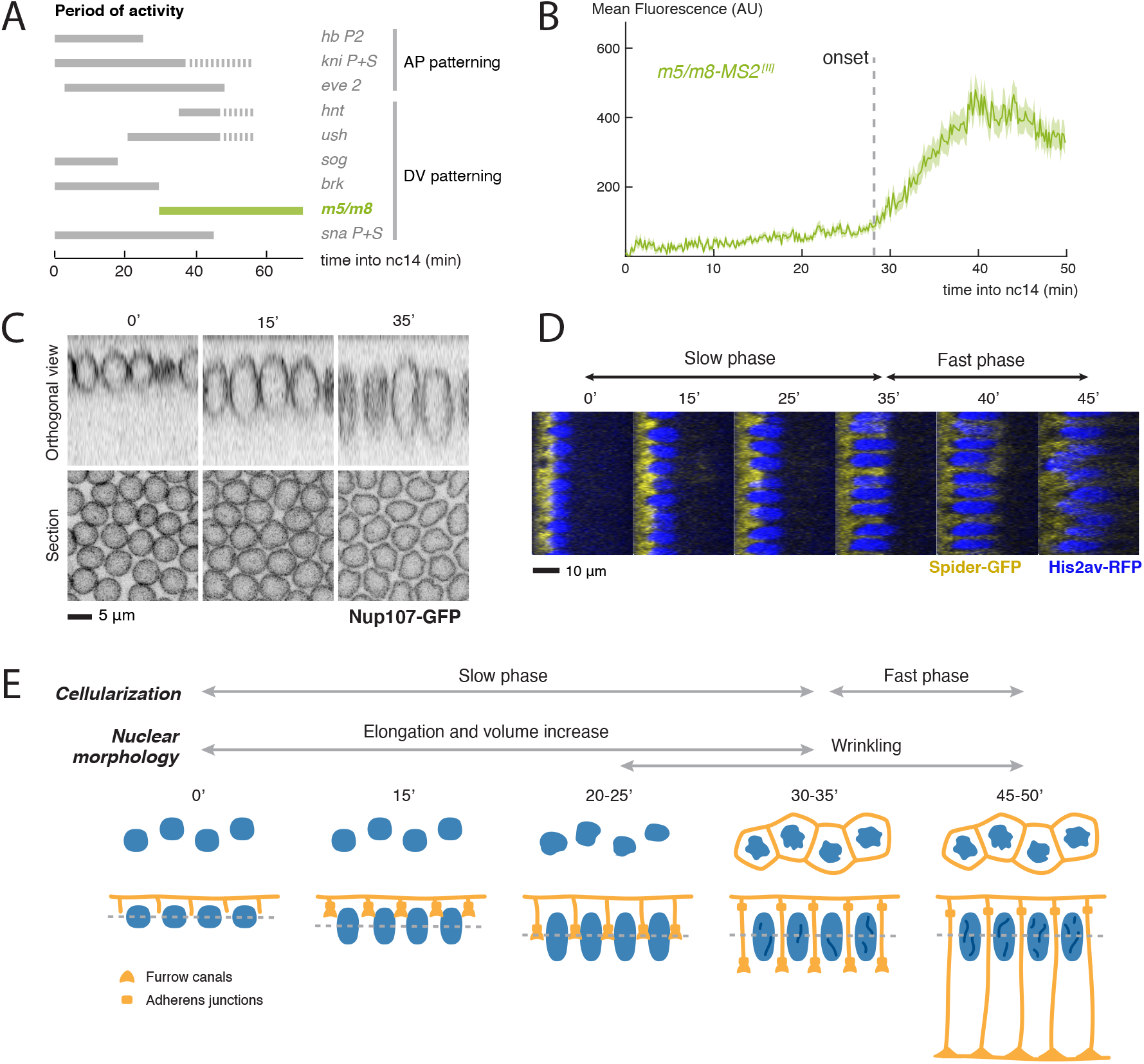
Correlation between developmental processes and onset of Notch dependent transcription. **A**) Summary of the expression timings of other published MS2 lines. Grey solid lines indicate activity and dashed lines indicate periods when transcription was not quantified but the enhancer/gene is expected to be active. Based on data from ***Garcia et al. 2013; Bothma et al. 2014; Bothma et al. 2015; Lim et al. 2017; Falo-Sanjuan et al. 2019; Hoppe et al. 2020*. B**) Mean profile of transcription from *m5/m8*. Transcription starts from 30 min into nc14. Based on data from ***Falo-Sanjuan et al. 2019*. C**) Medial section and orthogonal views of nc14 embryos at the indicated times (minutes into nc14) expressing the nuclear membrane marker Nup107-GFP. See also **Movie 1**. **D**) Orthogonal views of embryos expressing the cell membrane marker Spider-GFP and nuclei marker His2Av-RFP, indicating changes in nuclear and membrane length over time (minutes into nc14). **E**) Summary of the behaviours of nuclei and membranes during nc14. Cellularization takes place in two phases: slow (0-35’) and fast (35-55’) membrane in-growth. At the same time nuclei elongate and increase in volume (0-35’) and their surface becomes wrinkled from approximately 25 minutes onwards.

The substantial changes in nuclear morphology that occur during nc14 include shape-changes and alterations in pore clustering ***(Brandt et al. 2006; Pilot et al. 2006; Hampoelz et al. 2016)*.** To quantify these nuclear changes, embryos expressing Nup107-GFP ***(Katsani et al. 2008)*** were imaged live and the nuclear dimensions and eccentricity measured over time. Nuclei underwent substantial elongation in the apico-basal axis and increased in volume during the first 40 min of nc14 ***(Brandt et al. 2006; Pilot et al. 2006)*** (**Fig.** 1**C**, S1**A**, **Movie 1**). In addition, after approximately 25 min into nc14, there was an increase in eccentricity of nuclear medial slices, indicative of indentations (‘wrinkles’) appearing in the nuclear envelope (**Fig.** 1**C**, S1**A**, **Movie 1**) ***(Brandt et al. 2006; Pilot et al. 2006)*.** This transition to ‘wrinkling’ occurred around the time when signalling-dependent transcription is initiated.

Similarly, we used the membrane marker Spider-GFP (Gilgamesh, ***Morin et al. 2001***) to track the inward growing, lateral, membranes during cellularization and to quantify their growth. In agreement with previous reports, we could detect an initial slow phase of membrane ingrowth, which lasted circa 30-35 minutes, followed by a fast phase, which lasted circa 15 minutes and completed cellularization ***(Foe and Alberts 1983; Lecuit and Wieschaus 2000; Lecuit et al. 2002)*,** (**Fig.** 1**D**, S1**B**). By the end of the slow phase, membranes had reached the inferior margin of the nucleus (**Fig.** 1**D**). This corresponded approximately to the time at which Notch dependent transcription usually initiates ***(Falo-Sanjuan et al. 2019)*.** Furthermore, these ingrowing lateral membranes carried Notch and Delta. Tracking tagged full-length Notch (Notch-GFP, ***Couturier et al. 2012***) and endogenously-tagged Delta (Dl-mScarlet, ***Boukhatmi et al. 2020***) revealed that the location of both proteins expanded basally at the same rate as cellularization progressed (**Fig.** 2**A**, S2**A**) and that Dl-mScarlet tracked with E-cadherin (Shg-GFP) throughout cellularization (**Fig.** S2**BC**). Thus, lateral membranes containing Notch and Delta have partially formed at the time when signaling commences.

**Figure 2.**
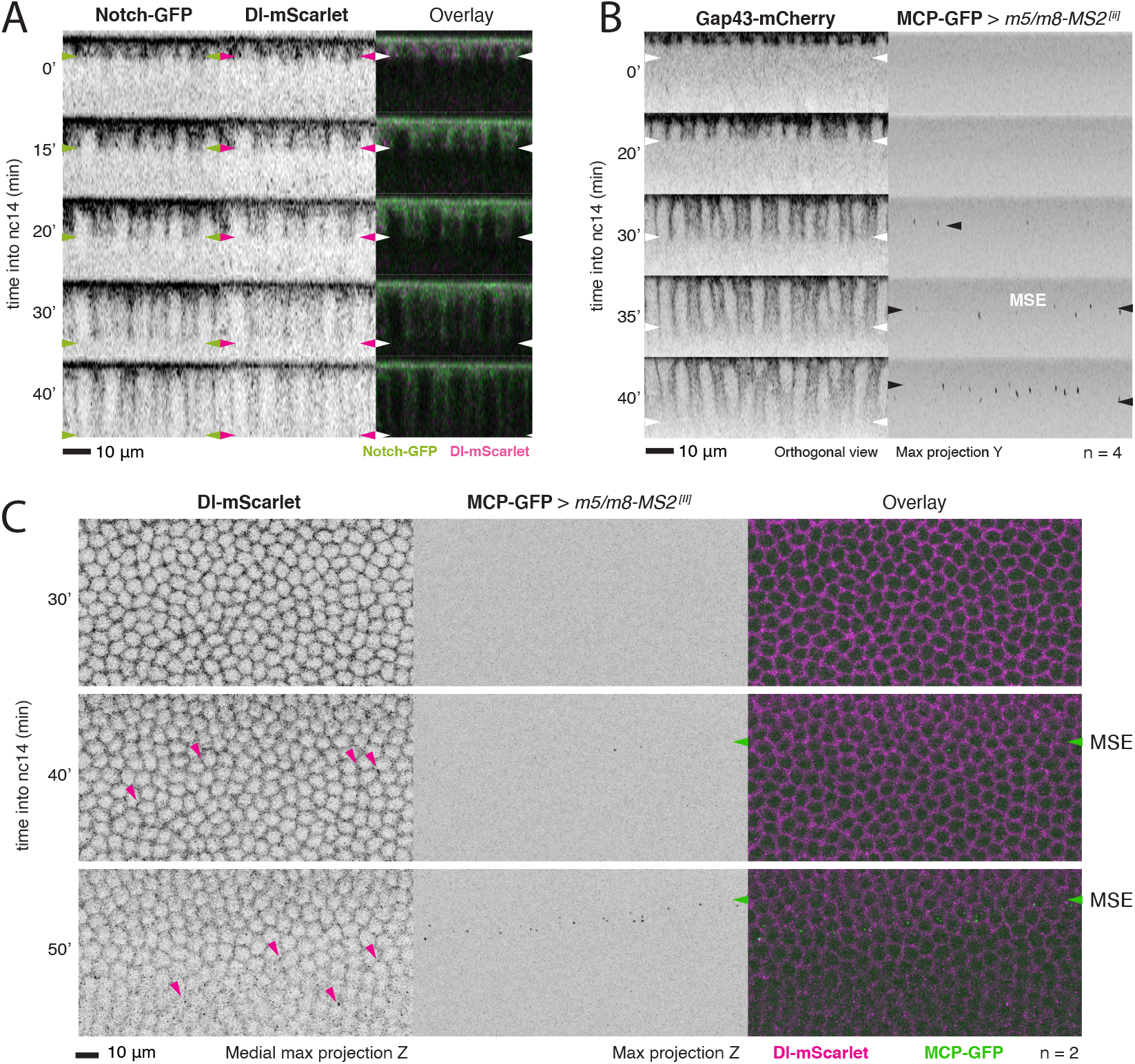
Notch responsive transcription starts before cellularization is completed. **A**) Orthogonal views of embryos expressing Notch-GFP and Dl-mScarlet, showing localization of Notch and Delta at cellularizing membranes. Arrowheads indicate position of the most basal accumulation. See also **Movie 2**. **B**) Stills of a movie of an embryo expressing the membrane marker Gap43-mCherry combined with MCP-GFP to image transcription from *m5/m8*. Orthogonal views of Gap43-mCherry (left) and maximum projections of the MCP-GFP channel (right) are shown. Time into nc14 (minutes) is indicated for each. Transcription starts from 30 min into nc14 and the whole mesectoderm stripe is visible by 35-40 min, before cellularization is completed. White arrowheads indicate position of cellularization front and black arrowheads indicate transcription in mesectoderm nuclei (MSE). See also **Movie 3**. **C**) Stills of an embryo expressing Dl-mScarlet combined with MCP-GFP to image transcription from *m5/m8*. Projections of medial slices (left, inverted image), maximum projections of the MCP-GFP channel (center, inverted image) and overlay of both (right) at three time-points as indicated. Delta can be detected in bright *puncta* (magenta arrowheads) close to the membrane in mesoderm cells from the time *m5/m8* transcription starts in the mesectoderm (MSE, green arrowheads).

To more precisely relate the time when Notch dependent transcription initiates with lateral membrane growth, we monitored transcription directed by the *m5/m8* enhancer in the presence of the membrane marker Gap43-mCherry ***(Fabrowski et al. 2013)*.** Results revealed that the onset of *m5/m8* dependent transcription occurred when the membrane had grown ~ 20 μm (**Fig.** 2**B**, **Movie 3**). At a similar stage, Delta membrane-levels became modulated in the mesodermal cells, where it was primarily detected in bright *puncta* close to the membrane. These changes occurred throughout the mesoderm, but not in the mesectodermal cells where *m5/m8* transcription was initiated (**Fig.** 2**C**), and likely correspond to increased Delta endocytosis driven by Neuralized, as reported previously ***(Morel et al. 2003; De Renzis et al. 2006)*.**

Based on the onset of the transcriptional read-out, these data indicate that productive Notch-Delta signaling is initiated after lateral membranes have started to form, during the transition between the slow and fast phases of membrane elongation, and significantly before cellularization finishes. This also corresponds to the period when the nuclei are undergoing morphological changes associated with the maturation of nuclear membranes and pores (**Fig.** 1**E**).

### Lateral membranes are limiting for Notch signalling

To distinguish the contributions from nuclear morphogenesis and lateral membrane formation on Notch signaling, we used mutations to perturb each process. First we investigated the consequences from disrupting nuclear shape-changes. *kugelkern* (*kuk*) encodes a nuclear lamina protein required for nuclear elongation and wrinkling at nc14 ***(Pilot et al. 2006; Brandt et al. 2006)*.** In *kuk[EY07696]* mutant embryos, although overall nuclear volume was unaffected, the nuclei had significantly reduced eccentricity, correlating with a reduction in their indentations (**Fig.** 3**AB**, S3**A**). Despite these modifications, transcription directed by *m5/m8* was unaltered; mean levels, onset and transcription profiles were similar to controls (**Fig.** 3**CDE**, S3**B**). These data suggest that the stage-specific changes in nuclear morphology are not required for the normal onset and levels of Notch dependent transcription.

**Figure 3.**
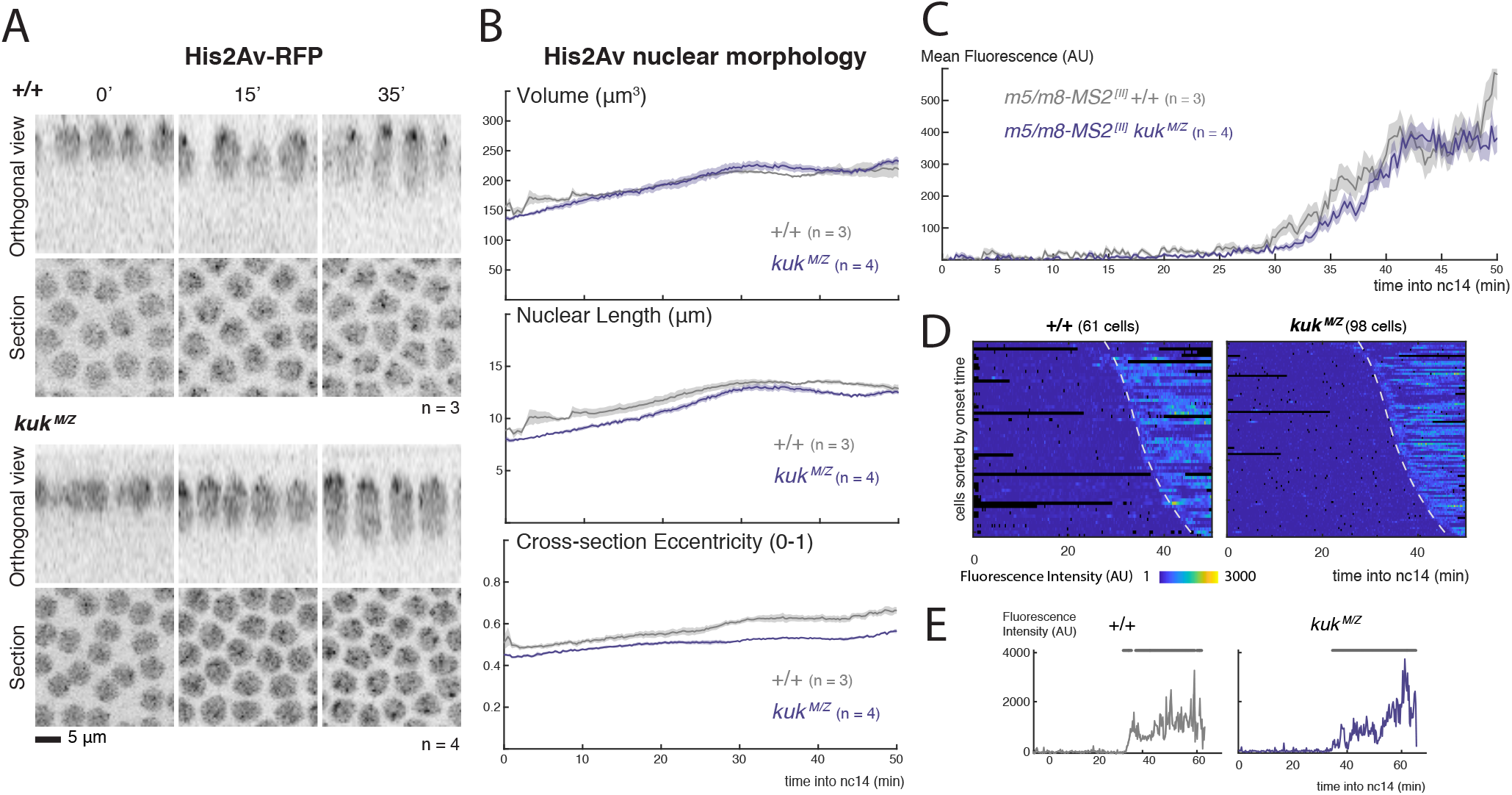
Changes in nuclear morphology do not influence Notch dependent transcription. **A)** Cross-sections and orthogonal views of the nuclear marker His2Av-RFP in wild type (top) and embryos obtained from homozygous *kuk* parents (*kuk^M/Z^*, bottom) at the indicated times (min into nc14). **B**) Quantification of nuclear morphological properties over time using the His2Av channel from MS2 experiments: volume, nuclear length and eccentricity of the medial slice. Mean and SEM (shaded area) of the mean properties calculated for each embryo is shown (n embryo numbers per condition indicated in each). **C**) Mean profiles of *m5/m8 ^[II]^* activity of mesectoderm nuclei in control and *kuk* embryos. Mean and SEM (shaded area) of all cells combined from multiple embryos are shown (n embryo numbers indicated in each). **D**) Heatmaps of transcription in all mesectoderm nuclei from sorted by onset time (top). Dashed lines indicate onset times in the control. **E**) Examples of transcription traces from mesectoderm nuclei. Grey lines indicate ON periods.

Second, we asked whether the formation of lateral membranes is a limiting factor in pathway activation, by analyzing the consequences on *m5/m8* transcription from a mutation in the zygotic gene *slam*, which disrupts cellularization ***(Lecuit et al. 2002)*.** Membrane formation was quantified by capturing a cross-section of the embryo in every time-point using transmitted light (**Fig.** 4**B**), and measuring the length of lateral membranes to determine the time-points when the cellularization front reached specific positions (**Fig.** 1**E**) ***(Lecuit and Wieschaus 2000; Lecuit et al. 2002)*.** As expected, homozygous mutant embryos for *slam* exhibited disrupted cellularization and severe morphological defects: all phases of cellularization were slowed down and, in 3 out of 4 cases, celluarization was not completed (**Fig.** 4**BC**). Strikingly, mesectoderm nuclei exhibited almost no transcriptional activity in *slam* mutant embryos. A few nuclei initiated some sporadic activity at the same time as in control embryos, but this lasted only few minutes (**Fig.** 4**EG**, S4**A**, **Movie 4**). As a result, only scattered nuclei were active in the mesectodermal stripe (**Fig.** 4**F**) and mean levels of transcription were close to background (**Fig.** 4**D**). These data argue that, in contrast to nuclear morphogenesis, normal lateral membrane formation is important for signalling to initiate and be maintained.

A proportion of the non-homozygous embryos displayed slowed, rather than blocked, cellularization (**Fig.** 4**C**). These were likely *slam^+/-^* heterozygotes, but could not be definitively distinguished from any pseudo-normal homozygous balancer embryos. Transcription onset and cellularization times were therefore quantified in all non *slam^-/-^* embryos in an unbiased way and plotted alongside controls. The results revealed a striking relationship between cellularization time-points and the onset of *m5/m8* activity (first quartile of onset times), with the strongest correlation when membranes reached the basal end of nuclei (*R*^2^ = 0.76) (**Fig.** 4**H**, S4**B**). Delta localization appeared normal in *slam* mutant embryos during the first 15-20 min of nc14 and then tracked with the delayed growth of the lateral membranes (**Fig.** S4**CD**). Delta thus occupies the available lateral membrane territory in each condition.

**Figure 4.**
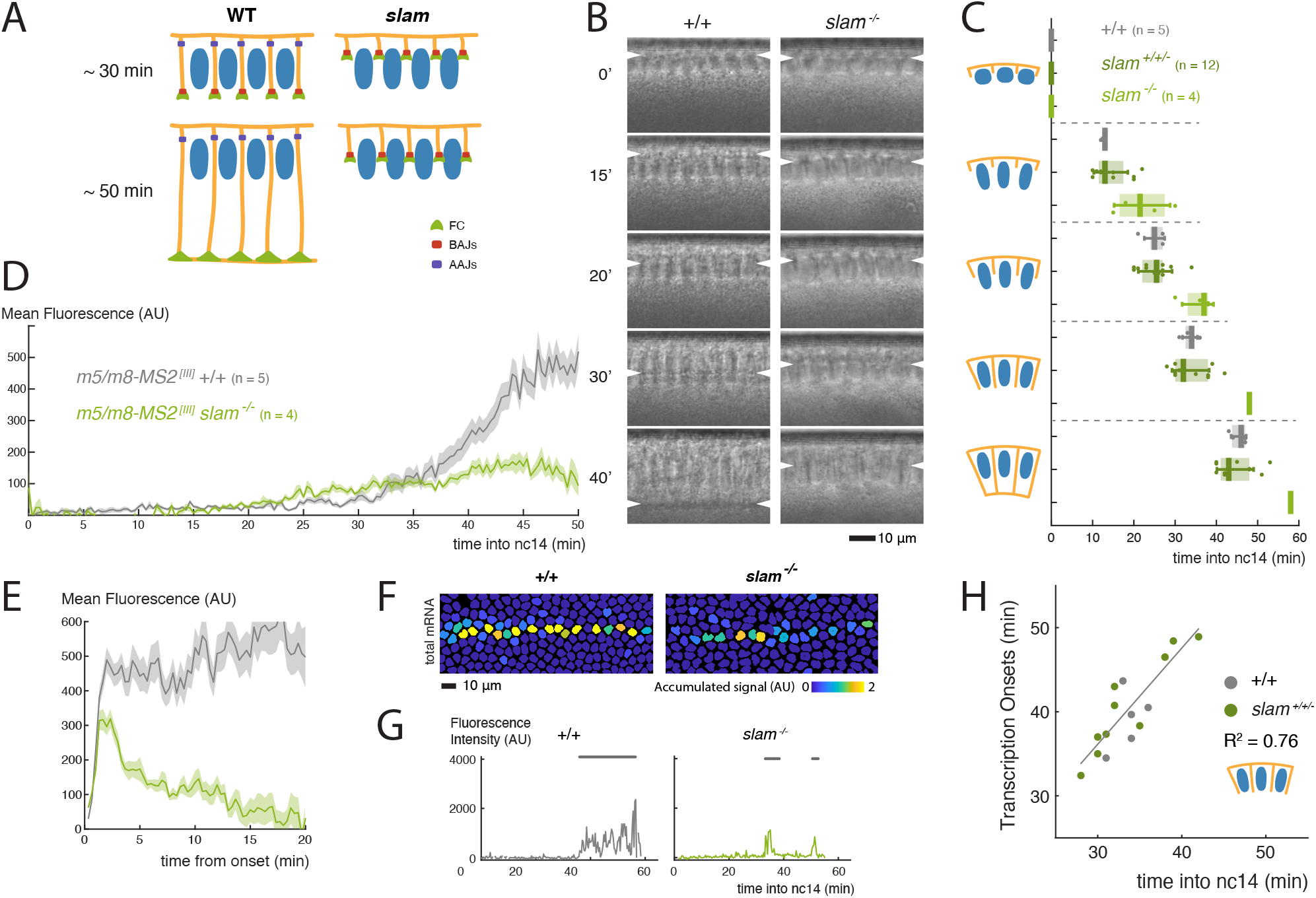
Lateral membranes are required for Notch signalling. **A**) Schematic representation of the effects on membrane formation produced by mutation in *slam*. **B**) Cross-sections of wild type and *slam^-/-^* embryos captured with transmitted light and used to measure cellularization progression. Arrowheads indicate position of the cellularization front. **C**) Boxplots indicating timing of cellularization progression (timepoints when membranes reach each of the lengths with respect to nuclei indicated in the cartoons) in wild type, *slam^+/+/-^* and *slam^-/-^* embryos. Median, Q1/Q3 quartiles and SD shown. **D**) Mean profile of *m5/m8 ^[III]^* activity in *slam^-/-^* embryos compared to controls. **E**) Mean levels of transcription in *slam^-/-^* embryos compared to controls when nuclei are aligned by their onset times. **F**) Tracked mesectoderm nuclei color-coded for their total transcription levels. **G**) Examples of transcription traces from mesectoderm nuclei. Grey lines indicate ON periods. **H**) Correlation between the timepoint of cellularization when membranes reach the basal end of nuclei with onset of transcription from *m5/m8* (calculated as the first quartile of onset times) in *slam^+/+/-^* and control embryos. R2 coefficients are calculated after pooling all points together. In **D** and **E**, mean and SEM (shaded area) of all cells combined from multiple embryos are shown (embryo numbers, n, indicated in C). See also **Movie 4**.

Together, these observations are consistent with Delta-Notch signalling initiating from the lateral membranes before cellularization finishes. The correlation between onset of transcription and membrane progression suggests that a specific step during cellularization determines when signalling can start. One possibility is that the membrane length *per se* is limiting because it determines the amount of Notch and Delta that are available for signaling. Alternatively, the formation of a specific membrane domain or junction may be the limiting factor that enables productive Notch-Delta interactions.

### Adherens junctions contribute to Notch activation

To investigate whether the onset of signaling is limited by the dimensions of the lateral membrane *per se* or by the establishment of specific domains, such as AJs, we first examined the transcriptional profiles in embryos mutant for *nullo*. Mutations in *nullo* destabilize furrow canals and mislocalize furrow canal components, with one consequence being that the transient basal adherens junctions (BAJs) do not form properly ***(Hunter and Wieschaus 2000; Hunter et al. 2002)*** (**Fig.** 5**A**) although apical adherens junctions (AAJs) are subsequently established. Unlike *slam* mutants, the cellularization front in *nullo* hemizygous embryos progressed at a similar mean rate to control embryos (**Fig.** 5**BC**). In accordance, mean transcription levels resembled those of control embryos (**Fig.** S5**ABC**, **Movie 4**), except for a small (<5 min) delay in onset times when all transcription profiles were considered together (**Fig.** 5**E**). However, on an embryo by embryo basis, onset times were more variable than controls and, strikingly, there was no correlation between lateral membrane length and transcription onset time in *nullo* hemizygous or heterozygous embryos (**Fig.** 5**F**, S5**D**), unlike with *slam* mutants. Overall, these observations suggest that lateral membrane length is not the limiting parameter for initiation of Notch signaling. Furthermore, as *nullo* mutations affect the initial assembly of BAJs, the fact that Notch dependent transcription is largely normal, argues that BAJ formation is not essential. We note however, that a few nuclei failed to initiate transcription. This is likely because some furrow canals fail to develop in *nullo* embryos. This loss of lateral membranes results in a disorganized and patchy stripe of mesectodermal transcription (**Fig.** 5**D**).

**Figure 5.**
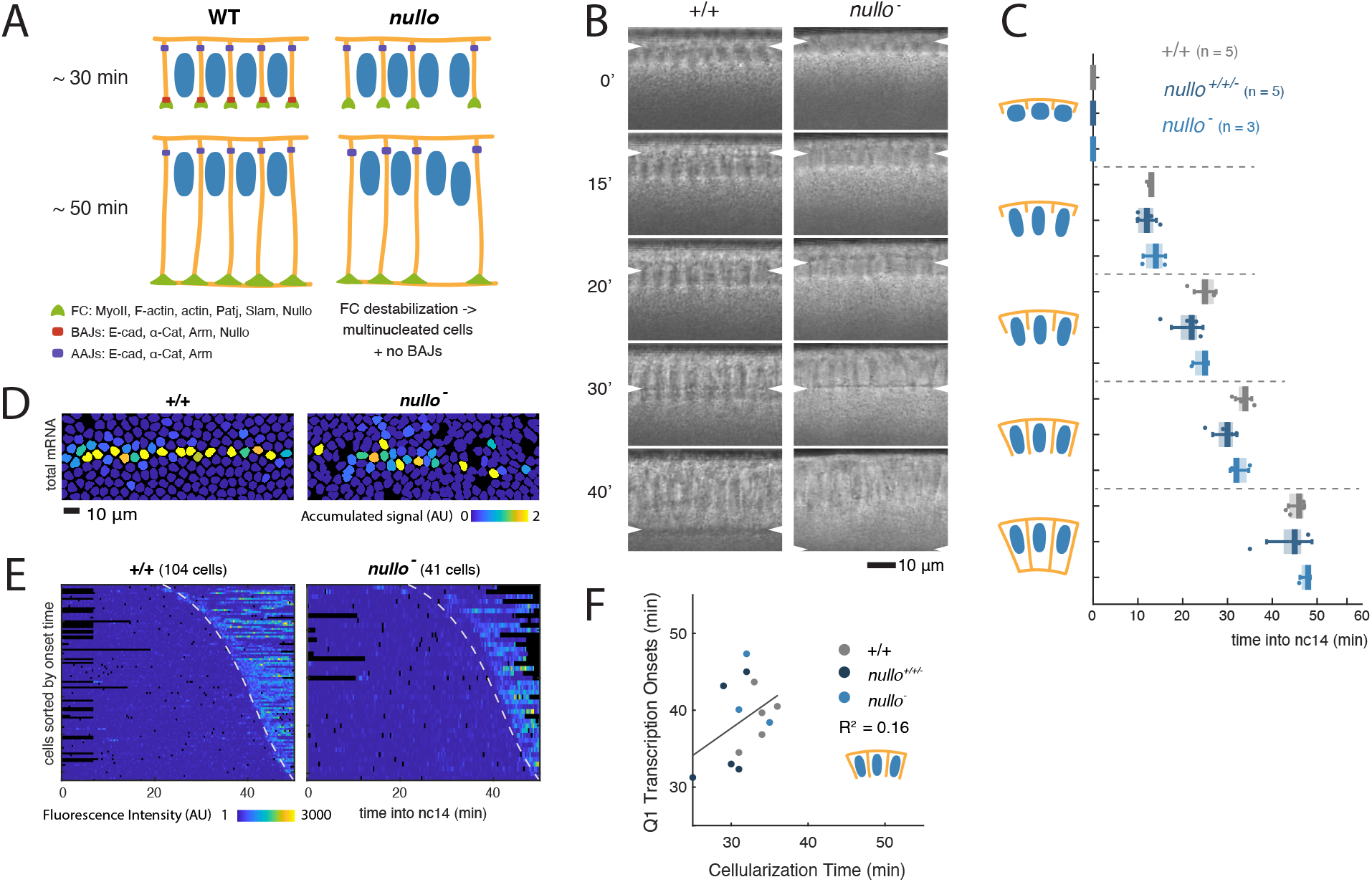
Defects in cellularization from absence of Nullo perturb Notch signalling independently of membrane growth. **A**) Schematic representation of the effects on membrane formation produced by mutations in *nullo*. **B**) Cross-sections of wild type and *nullo^−^* embryos captured with transmitted light and used to measure cellularization progression. Arrowheads indicate position of cellularization front. **C**) Boxplots indicating timing of cellularization progression (timepoints when membranes reach each of the lengths with respect to nuclei indicated in the cartoons) in wild type, *nullo^+/+/-^* and *nullo^−^* embryos. Median, Q1/Q3 quartiles and SD shown. **D**) Tracked mesectoderm nuclei color-coded for their total transcription levels. **E**) Heatmaps of *m5/m8 ^[III]^* transcription in all mesectoderm nuclei sorted by onset time. Dashed lines indicate onset times in wild type. **F**) Correlation between the timepoint of cellularization when membranes reach the basal end of nuclei with onset of transcription from *m5/m8 ^[III]^* (calculated as the first quartile of onset times) in *nullo^+/+/-^* and control embryos. R2 coefficients are calculated after pooling all points together. Images, plots and quantifications of control embryos are duplicated from **Fig.** 4. See also **Movie 4**.

Although *nullo* mutants lack BAJs, the formation of AAJs appears normal ***(Hunter and Wieschaus 2000)*.** We next investigated the consequences on *m5/m8* directed transcription of disrupting both basal and apical AJs, by depleting the key junctional linker, α-Catenin (α-Cat) ***(Staller et al. 2013).*** Maternal RNAi knockdown (KD) led to a marked depletion of α*-Cat* mRNA and protein (**Fig.** S6**AB**), resulting in 100% embryos with gastrulation failure but with only modest delays in cellularization (**Fig.** S6**C**). Strikingly, Notch dependent transcription was affected in these α-Cat KD embryos in advance of any gastrulation defects. The main consequences were a disruption of the mesectodermal stripe (**Fig.** 6**A**) and an overall reduction in the mean levels of transcription without affecting the onset times (**Fig.** 6**CD**, **Movie 5**). This was due to a shift in the distribution of activity-levels, with many nuclei exhibiting a marked reduction in their overall mRNA output (**Fig.** 6**E**, S6**D**). To determine whether α-Cat contribution to Notch signalling is relevant in the context of endogenous gene activity, we tagged with MS2 loops one of the Notch target genes proposed to be regulated by the *m5/m8* enhancer - *E(spl)m8-HLH* ***(Zinzen et al. 2006)*.** In a similar way to *m5/m8 ^[III]^*, *E(spl)m8-HLH* transcription was disrupted upon α-Cat KD: the mesectodermal stripe was disorganized, the mean levels were reduced without a change in onset times, and the range of accumulated mRNA levels per nucleus was diminished (**Fig.** 6**BFGH**, S6**E**, **Movie 6**). Overall, these results suggest that the formation of AJs is an important step in the timing and strength of Notch activation during nc14. When perturbed, reduced levels of Notch dependent transcription occurred.

**Figure 6.**
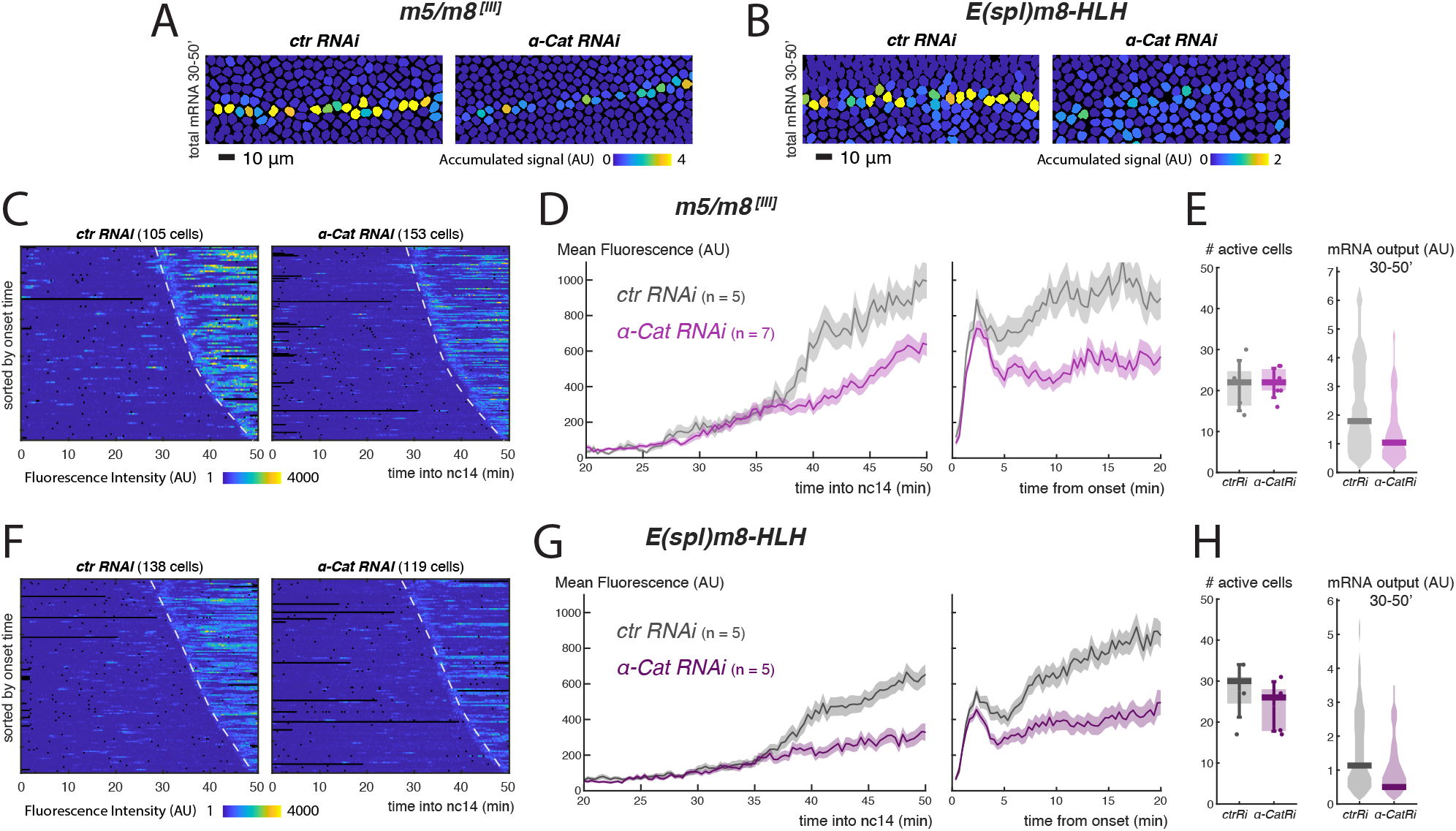
Adherens junctions influence Notch dependent transcription. **A**-**B**) Tracked mesectoderm nuclei color-coded for total *m5/m8 ^[III]^* (**A**) or *E(spl)m8-HLH* (**B**) transcription (accumulated signal from 30 to 50 min into nc14). **C**) Heatmaps of *m5/m8 ^[III]^* transcription in all mesectoderm nuclei sorted by onset time in control and α-Cat depleted embryos. Dashed lines indicate onset times in controls. **D**) Mean profile of *m5/m8 ^[III]^* activity in α*-Cat RNAi* embryos compared to controls, aligned by developmental time (left) or transcription onset times (right). **E**) Boxplot indicating number of mesectoderm cells transcribing *m5/m8 ^[III]^* in each embryo in control and α*-Cat RNAi* embryos (left, median, Q1/Q3 quartiles and SD shown) and violin plot showing the distribution of output levels of transcription (accumulated signal from 30 to 50 min into nc14, as in **A**) combining all nuclei from each condition (right, distribution and median shown). **F**) Heatmaps of *E(spl)m8-HLH* transcription in all mesectoderm nuclei sorted by onset time in control and α-Cat depleted embryos. Dashed lines indicate onset times in controls. **G**) Mean profile of *E(spl)m8-HLH* activity in α*-Cat RNAi* embryos compared to controls, aligned by developmental time (left) or transcription onset times (right). **H**) Boxplot indicating number of mesectoderm cells transcribing *E(spl)m8-HLH* in each embryo in control and α*-Cat RNAi* embryos (left, median, Q1/Q3 quartiles and SD shown) and violin plot showing the distribution of output levels of transcription (accumulated signal from 30 to 50 min into nc14, as in **B**) combining all nuclei from each condition (right, distribution and median shown). In **D** and **G**, mean and SEM (shaded area) of all cells combined from multiple embryos are shown (n embryo numbers indicated in each). See also **Movie 5** and **Movie 6**.

To investigate whether the role of α-Catenin and AJs was likely to involve direct effects on Notch, we used SIM (Structured Illumination microscopy) to assess the extent of protein co-localization. The high resolution imaging revealed a heterogenous distribution of Notch along the growing lateral membranes. Apically, Notch levels were similar around the whole circumference, whereas sub-apically Notch was enriched at tricellular junctions and more basally it was present in the furrow canals (the most basal part of growing membranes), delineated by F-actin (**Fig.** 7**A**). E-cadherin was also detected in all these positions, but the two proteins were distributed unevenly in membrane clusters with relatively few sites where they were co-localized (**Fig.** 7**A**). Overall, the low level of co-localization suggests that Notch is not directly sequestered into the AJs, although it is in close proximity. Furthermore, Notch localization was not disrupted upon α-Catenin depletion. In embryos at mid-cellularization (around the time Notch dependent transcription initiates), Notch was present at a similar level and with similar overall distribution as in α-Catenin depleted embryos (**Fig.** 7**B**, S7**B**). Although defects in adhesion became evident at late cellularization, in the form of “holes” at the tricellular junctions ***(Yu and Zallen 2020)*** that also displaced Notch into a surrounding ring (**Fig.** S7**A**), no other changes in Notch localization were apparent, leading us to conclude that α-Catenin depletion does not generally disrupt the distribution of Notch in the lateral membranes, despite its effect on Notch dependent transcription.

**Figure 7.**
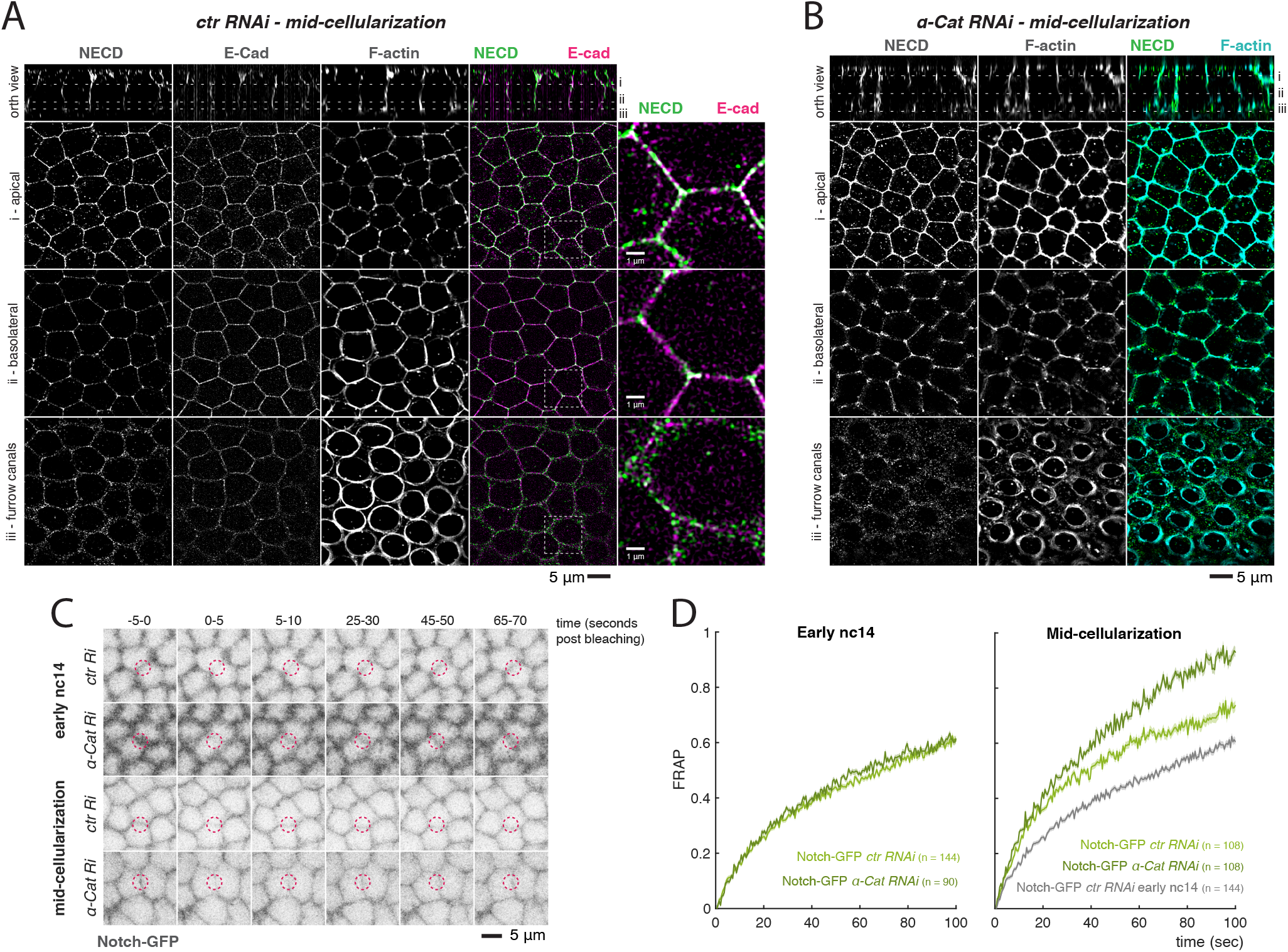
α-Catenin depletion influences Notch membrane dynamics but not localization. **A**-**B**) Mid-cellularization control (**A**) or α*-Cat* (**B**) RNAi embryos stained with phalloidin and antibodies against the extracellular domain if Notch (NECD) and E-cad and imaged using SIM (E-cad channel not shown in **B**). Top panels are orthogonal views with lines marking individual planes shown below. **C**) Stills of Notch-GFP FRAP experiments in the indicated conditions in early nc14 or mid-cellularization embryos. Each still is an average of 10 frames between the indicated timepoints. Red circles indicate the quantified region over time. **D**) FRAP experiments performed on Notch-GFP in early nc14 (left) and mid-cellularization (right) embryos, comparing control and α-Cat depletion.

α-Catenin is proposed to influence E-cad stability at the membrane ***(Bajpai et al. 2008; Jurado et al. 2016; Ishiyama et al. 2018)*.** We thus wondered if α-Catenin depletion could similarly be influencing Notch stability, rather than localization. To this end, we measured the fluorescence recovery after photobleaching (FRAP) of Notch-GFP ***(Couturier et al. 2012)*,** as an indication of its turnover in the membrane. There was a notable change in the speed of recovery between early nc14 and mid-cellularization time-points, with faster recoveries detected at the later time-point, suggesting there is more rapid turnover of Notch in the membrane around the time that signalling commences (**Fig.** 7**CD**). However, as the measurements were made at random locations in the embryo, the differences represent general properties of Notch at this time, rather than any signaling induced changes, as the latter would be restricted to mesectodermal cells. α-Catenin depletion had no effect on the Notch recovery at the early time point. However, at mid-cellularization, α-Catenin depletion resulted in faster recovery times (**Fig.** 7**CD**), suggesting that it normally restricts the turnover or recycling. One consequence from the α-Catenin dependent effects on membrane stability is that Notch would have a longer residence time in the membrane, which could permit higher levels

## Discussion

The geometry of a tissue and the nature of the cell contacts are likely to be important factors influencing the levels and duration of Notch signalling ***(Shaya et al. 2017)*.** By analyzing the transcriptional output of Notch signalling in live blastoderm embryos we have been able to relate the time of productive ligand-receptor interactions with landmarks in cellular membrane growth. Strikingly, signalling was initiated after lateral membranes had grown to approximately 1/3 of their final length but before cellularization was complete. There was a strong correlation between cellularization time in each embryo, measured by the length of the lateral membranes, and onset of transcription, even in embryos where membrane growth was delayed. These results argue that a key step during membrane morphogenesis determines when signalling can initiate. The same restrictions could also influence when signalling can re-initiate following cell division.

The requirement for lateral membrane growth and morphogenesis can explain why two different Notch-responsive enhancers initiate transcription within a few minutes of each other ***(Falo-Sanjuan et al. 2019)*,** because there would be a coordinated release of NICD when the receptor and ligands first became juxtaposed. Furthermore, the correlations, together with the lack of transcription in *slam* mutant embryos, are hard to reconcile with the model that NICD accumulates in the nucleus from the beginning of nc14 ***(Viswanathan et al. 2019)*.** Our results also favour the model that signaling is initiated *in trans*, between receptor and ligand located on neighbouring cell membranes, rather than *in cis*, between ligand and receptors on the same apical and/or internal membranes ***(Coumailleau et al. 2009; Nandagopal et al. 2019)*.**

One plausible explanation for the precise onset of transcription at a specific moment during membrane morphogenesis could be that a minimal area of interface is required for signalling to surpass a critical threshold. However, our data argue against this being the limiting factor and suggest that the formation and/or maturation of membrane domains or junctions is required. First, a delay in transcription onsets was observed in *nullo* mutants, despite normal cellularization speed, where the transcription onset and lateral membrane growth were no longer strongly correlated. Second, Notch responsive transcription was impaired when α-Cat, a key component of AJs, was depleted. The number of nuclei with high levels of transcription from the *m5/m8* enhancer was reduced, leading to a reduction in the overall mean levels. Similar effects on the endogenous *E(spl)m8-HLH* were also seen upon α-Cat depletion. As the lateral membranes are fully formed in these depleted embryos, the results suggest that features coordinated by AJs are important for normal signalling. Given the variability of the effects on transcription, it is likely that these properties are required to achieve high levels of Notch signalling rather than being absolutely required for Notch activation.

The effects of AJs on Notch signalling could be direct or indirect. Based on superresolution imaging, there was no specific co-enrichment of Notch with components of AJs, such as cadherin, nor was Notch localization adversely affected by α-Cat depletion. Together, these results make it unlikely that the direct recruitment of Notch to apical junctions is a limiting factor. However, Notch dynamics at the membrane were altered in α-Cat depleted embryos, based on FRAP experiments. These indicated that the membrane associated Notch is less stable when α-Cat is depleted, which could reduce the amount of Notch that is available to interact and signal at any one moment ***(Khait et al. 2016).*** It is not possible to distinguish whether the altered dynamics are due to changes in recycling/synthesis or in lateral diffusion. As the latter could also result in altered segregation of Notch and the g-secretase cleavage machinery ***(Kwak et al. 2020)*,** either change could explain the reduced transcription output in the α-Cat depleted embryos. An alternative explanation is that α-Cat, and AJs, contribute to Notch activation because of their important role in cell-cell adhesion. α-Cat functions as the linker between AJs and actomyosin, and is involved in transmitting contractile forces across cells ***(Jurado et al. 2016)*.** AJ-mediated adhesion could promote higher Dlpulling force, hence enhancing Notch cleavage and NICD release ***(Gordon et al. 2015)*** to regulate outputs. It is also possible that α-Cat exerts its effects via a combination of mechanisms.

Our data that lateral membranes are required for signalling are consistent with elegant experiments tracking photoconverted receptor populations in *Drosophila* sensory organ precursors (SOP), which indicated that the lateral pool of Notch is the one that becomes activated ***(Trylinski et al. 2017)*.** In this context, the active receptor population was located basal to the apical junctions. In contrast, during vertebrate neurogenesis adherens junctions at the apical luminal surface of the neuronal progenitors have been proposed as the site of signaling ***(Hatakeyama et al. 2014)*.** As Notch does not strongly colocalize with Cadherin at cellularization, our results fit better with those from SOPs and from cell culture studies proposing that full length Notch is excluded from AJs ***(Kwak et al. 2020)*.** However, ligand interactions and postactivation cleavage may occur at different sites in the membrane and indeed the sites of ligand interactions may differ according to the tissue architecture. For example, in the *Drosophila* follicular epithelium, cells receive signals from the neighbouring germ cells via their apical surface ***(López-Schier and St Johnston 2001).*** In other contexts, basal actin-based protrusions and cytonemes have been proposed as a source of ligand mediating longer range signalling ***(Huang and Kornberg 2015; Hunter et al. 2019; Boukhatmi et al. 2020)*.** Nevertheless, it is evident from the results presented here that the cell architecture, and the formation of apical junctions, are important features in enabling signalling in a simple epithelium. It will be interesting to see in which other contexts adherens junctions contribute to Notch activity. For example, a recent study showed AJs disruption in the mouse brain led to a phenotype of early differentiation of progenitor cells similar to that caused by reduced Notch signalling ***(Kwak et al. 2020)*,** suggesting there might be a widespread role of adherens junction in modulating Notch activity.

## Methods

### Fly strains and genetics

The following *Drosophila* strains were used: *sqh-Gap43-mCherry* ***(Izquierdo et al. 2018)*,** *GFP-gish[Spider]* (BDSC #59025, ***Morin et al. 2001***), *shg-GFP* (BDSC #60584, ***Huang et al. 2009***), *Notch-GFP* (*Ni-GFP* from ***Couturier et al. 2012***), *Dl-mScarlet* ***(Boukhatmi et al. 2020)*,** *Nup107-GFP* (BDSC #35514, ***Katsani et al. 2008***), *nos-MCP-GFP* (II, BDSC #63821) and *His2Av-RFP*; *nos-MCP-GFP* (BDSC #60340, ***Garcia et al. 2013***), *His2Av-RFP* (III, BDSC #23650). The *m5/m8-peve-MS2-lacZ* second chromosome (*m5/m8 ^[II]^*) and third chromosome (*m5/m8 ^[III]^*) MS2 reporter lines were generated in ***(Falo-Sanjuan et al. 2019)*.** *E(spl)m8-HLH-MS2* was generated during this work. Full genotypes of used lines are detailed in **Table** 1.

**Table 1.**
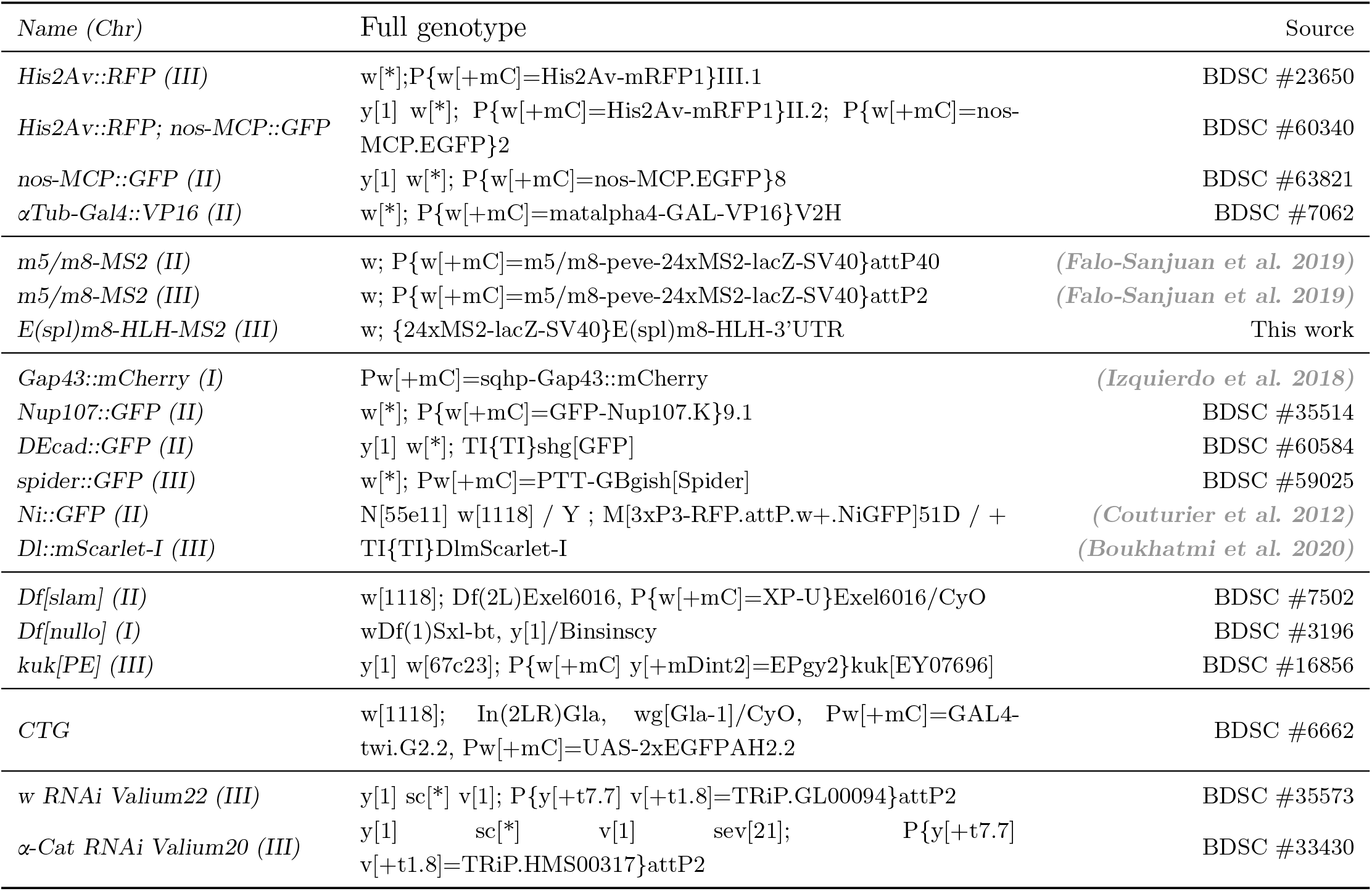
Full genotypes of used *Drosophila* lines.

#### Generation of endogenously tagged E(spl)m8-HLH-MS2

24 MS2 loops, lacZ and SV40 (5.4 kb in total, same as used for the *m5/m8* reporter) were inserted in the genome by CRISPR/Cas9 scarless genome engineering (flycrispr.org) to replace the *E(spl)m8-HLH-MS2* 3’UTR while keeping its coding sequence intact. Briefly, a plasmid containing homology arms flanking *E(spl)m8-HLH-MS2* 3’UTR, lacZ, SV40 and the PiggyBac 3xPax3-dsRED cassette from *pHD-ScarlessDsRed* (flycrispr.org) was synthesized by NBS Biologicals (Huntingdon, England). 24 MS2 loops from *pCR4-24XMS2SL-stable* (Addgene #31865) were subsequently inserted using an *EcoR*I site. Transformants were obtained by co-injecting (performed by the Genetics Fly Facility, University of Cambridge) this plasmid with a pCFD3-dU6:3gRNA plasmid (Addgene #49410) expressing the gRNA *CTGTGATAGCCCAACTGTGA* and screening for 3xPax3-dsRED. The *3xPax3-dsRED* cassette was excised by crossing with aTub84B-PiggyBac flies (BDSC #32070). Maps of the homology and gRNA plasmids and final genomic sequence have been deposited at: https://benchling.com/braylab/f/tE0Fz0Q1-endogenous-ms2-lines/.

#### Mutant backgrounds

To test expression from *m5/m8* in the *kuk[PE]* mutant background, a second chromosome recombinant *His2av-RFP, nos-MCP-GFP* ***(Falo-Sanjuan et al. 2019)*** was combined with *kuk[EY07696]* (BDSC #16856, ***Pilot et al. 2006***). *m5/m8 ^[II]^* was also combined with *kuk[EY07696]* and, since *kuk[EY07696]* is homozygous viable, *His2av-RFP, nos-MCP-GFP* / CyO; *kuk[EY07696]* females were crossed with *m5/m8 ^[II]^*; *kuk[EY07696]* males to obtain embryos that were maternal and zygotic mutant for this hypomorphic *kuk* allele. Control embryos were obtained by crossing *His2av-RFP, nos-MCP-GFP* / CyO females with *m5/m8 ^[II]^* males.

To test expression from *m5/m8* in the *slam* and *nullo* mutant backgrounds, third chromosome recombinants *His2av-RFP, nos-MCP-GFP* ***(Falo-Sanjuan et al. 2019)*** were combined with deficiencies encompassing *nullo* (*Df(1)Sxl-bt*, BDSC #3196) or *slam* (*Df(2L)Exel6016, Pw[+mC]=XP-UExel6016t*, BDSC #7502). *m5/m8 ^[III]^* was also combined with *Df[slam]*. Control embryos were obtained by crossing *His2av-RFP, nos-MCP-GFP* females with *m5/m8 ^[III]^* males. Homozygous mutant embryos for *slam* were obtained from crossing *Df[slam]* / *CTG*; *His2av-RFP, nos-MCP-GFP* with *Df[slam]* / *CTG*; *m5/m8 ^[III]^* and they were recognized by the absence of the labelled chromosome *CTG* (*CyO-twi-GFP*, BDSC #6662). Hemizygous embryos for *nullo* were obtained from crossing *Df[nullo]* / *FM6*;; *His2av-RFP, nos-MCP-GFP* with *m5/m8 ^[III]^* and they were recognized by defects in cellularization and gastrulation and lethality. In the two tested genes, 1 out of 4 embryos are expected to be mutant or the desired gene. The other embryos obtained from the same cross were grouped together as controls and to test a potential effect of heterozygous mutants. Note in this group 2/3 of the embryos are heterozygous for each tested gene and 1/3 contained 2 wild type alleles, though in some cases contained in balancer chromosomes.

#### Maternal KD

The maternal driver α*Tub-Gal4::VP16* (BDSC # 7062) was combined with *His2av-RFP, nos-MCP-GFP* to generate a*Tub-Gal4::VP16*; *His2Av-RFP*, *nos-MCP-GFP*. To knock down α*-Cat* from the maternal germline this stock was crossed with *UASp-*α*-Cat-RNAi* (BDSC #33430) or *UASp-w-RNAi* as control (BDSC #35573) and females a*Tub-Gal4::VP16* / + ; *His2Av-RFP*, *nos-MCP-GFP* / *UASp-RNAi* were crossed with *m5/m8 ^[III]^* to obtain the experimental embryos. To quantify the degree of maternal KD, a*Tub-Gal4::VP16* was crossed with the same lines and F2 embryos were collected for antibody staining and RT-qPCR.

Crosses used for each experiment are detailed in **Table** 2.

**Table 2.** Genotypes used in each experiment.

| Cross | Figure |
| --- | --- |
| ♀ *His2Av::RFP* ; *nos-MCP::GFP* x ♂ *m5/m8-peve-MS2-lacZ-SV40*[attP40,II] | 1B |
| ♀ x ♂ *Nup107::GFP* | 1C, S1A, S3A |
| ♀ x ♂ *Spider::GFP* ; *His2Av::RFP* | 1B |
| ♀ x ♂ *Ni::GFP* / + ; *Dl::mScarlet* / + | 2A, S2A |
| ♀ *Gap43::mCherry* ;; *nos-MCP::GFP* x ♂ *m5/m8-peve-MS2-lacZ-SV40*[attP40,II] | 2B |
| ♀ x ♂ *nos-MCP::GFP* / *m5/m8-peve-MS2-lacZ-SV40*[attP40,II] ; *Dl::mScarlet* / + | 2C |
| ♀ x ♂ *ECad::GFP* / + ; *Dl::mScarlet* / + | S2B, C |
| ♀ *His2Av::RFP*, *nos-MCP::GFP* / *CyO* x ♂ *m5/m8-peve-MS2-lacZ-SV40*[attP40,II] | 3, S3B |
| ♀ *His2Av::RFP*, *nos-MCP::GFP* / *CyO* ; *kuk[PE]*<br>x ♂ *m5/m8-peve-MS2-lacZ-SV40*[attP40,II] ; *kuk[PE]* | 3, S3B |
| ♀ x ♂ *Nup107::GFP* ; *kuk[PE]* / TM6B | S3A |
| ♀ *His2Av::RFP*, *nos-MCP::GFP* (III) x ♂ *m5/m8-peve-MS2-lacZ-SV40*[attP2,III] | 4, S4AB, 5, S5 |
| ♀ *Df[slam]* / CTG ; *His2Av::RFP*, *nos-MCP::GFP*<br>x ♂ *Df[slam]* / CTG ; *m5/m8-peve-MS2-lacZ-SV40*[attP2,III] | 4, S4AB |
| ♀ x ♂ *Df[slam]* / CTG ; *Dl-mScarlet* / TM6B | S4CD |
| ♀ *Df[nullo]* / FM6 ;; *His2Av::RFP*, *nos-MCP::GFP*<br>x ♂ *m5/m8-peve-MS2-lacZ-SV40*[attP2,III] | 5, S5 |
| ♀ *αTub-VP16* / + ; *His2Av::RFP*, *nos-MCP::GFP* / *UASp-w RNAi*<br>x ♂ *m5/m8-peve-MS2-lacZ-SV40*[attP2,III] | 6ACDE, S6CD |
| ♀ *αTub-VP16* / + ; *His2Av::RFP*, *nos-MCP::GFP* / *UASp-α-Cat RNAi*<br>x ♂ *m5/m8-peve-MS2-lacZ-SV40*[attP2,III] | 6ACDE, S6CD |
| ♀ *αTub-VP16* / + ; *His2Av::RFP*, *nos-MCP::GFP* / *UASp-w RNAi*<br>x ♂ *MS2-lacZ-SV40*[E(spl)m8-HLH-3'UTR] | 6BFGH, S6E |
| ♀ *αTub-VP16* / + ; *His2Av::RFP*, *nos-MCP::GFP* / *UASp-α-Cat RNAi*<br>x ♂ *MS2-lacZ-SV40*[E(spl)m8-HLH-3'UTR] | 6BFGH, S6E |
| ♀ x ♂ *αTub-VP16* / + ; *UASp-w RNAi* / + | S6AB, 7A |
| ♀ x ♂ *αTub-VP16* / + ; *UASp-α-Cat RNAi* / + | S6AB, 7B, S7A |
| ♀ x ♂ *αTub-VP16* / *Ni::GFP* ; *Dl::mScarlet* / *UASp-w RNAi* | 7DC, S7B |
| ♀ x ♂ *αTub-VP16* / *Ni::GFP* ; *Dl::mScarlet* / *UASp-α-Cat RNAi* | 7DC, S7B |

### Antibody staining

Embryos where dechorionated in bleach and fixed in a 1:1 mixture of heptane and 40% formaldehyde for 9 minutes. Embryos were then stuck to tape, manually devetillinized in PBS, and transfered to eppendorf tubes. Stainings to quantify maternal KD and for SIM were carried out in the same way: embryos were blocked in 1% BSA for 1h, incubated with primary antibodies overnight at 4C, washed in PBS-TritonX 0.1%, incubated with secondary antibodies for 2h at RT, washed in PBS-TritonX and mounted in Vectashield mounting medium. Used primary antibodies were: 1:100 concentrated rat DCAT-1 from DSHB (Developmental Studies Hybridoma Bank - Iowa, USA), 1:10 supernatant mouse NECD from DSHB C458-2H, 1:10 supernatant rat DCAD2 from DSHB. Used secondary antibodies were: 1:200 anti-Rat-FITC (Jackson Immunoresearch) for α-Cat KD quantification; 1:200 anti-Mouse-Alexa488 (Invitrogen), 1:200 anti-Rat-Alexa568 (ThermoFisher) for SIM. Embryos were also stained with 1:1500 Phalloidin-iFluor647 (Abcam).

### mRNA extraction and qPCR

Embryos were dechorionated in bleach and early embryos (pre-nc10) / eggs were selected in Voltalef medium. Pools of 15-20 embryos of each genotype were transfered to eppendorf tubes and dissociated in TRI Reagent (Sigma) with a plastic pestle. mRNA was extracted by adding chloroform, 10 min centrifugation at 4C and let to precipitate with isopropanol overnight. DNA was then pelleted by 10 min centrifugation at 4C, washed in 70% ethanol, dried and resuspended in DEPC-treated water. Approximately 2 μg of RNA from each sample were DNAse treated with the DNA-free™ DNA Removal Kit (Invitrogen) in the presence of RiboLock RNase Inhibitor (Thermo Scientific). 1 μg of DNA-free RNA was then used for reverse transcription using M-MLV Reverse Transcriptase (Promega) in the presence of RiboLock. Samples were diluted 1:2 for RT-qPCR using SYBR Green Mastermix (Sigma) and primers detailed in **Table** 3.

**Table 3.**
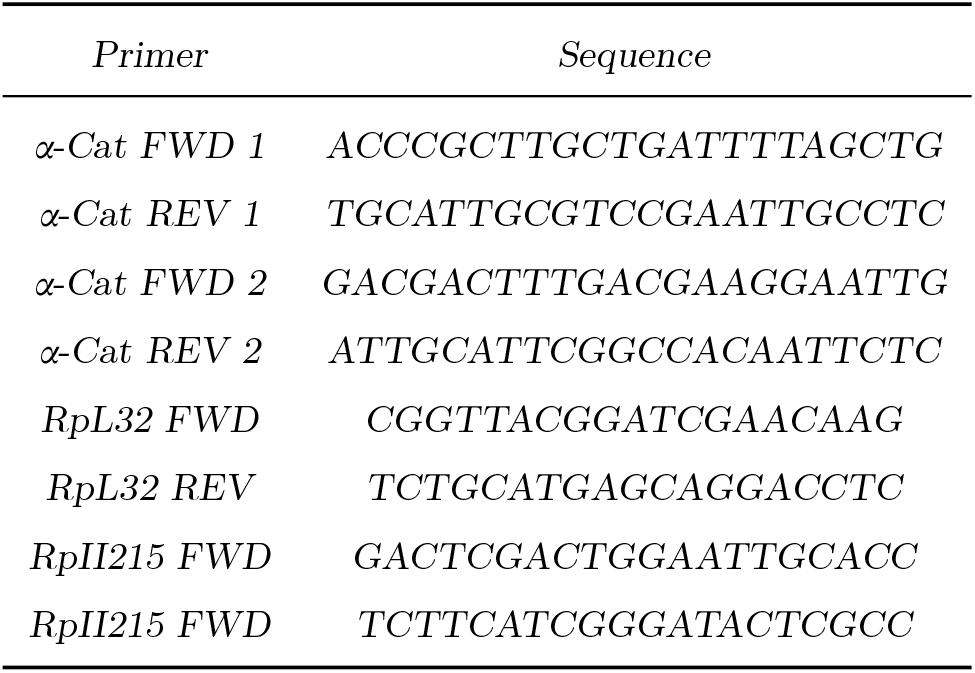
Primers used for qPCR.

### Structured Illumination Microscopy

Structured Illumination Microscopy (SIM) was carried out in stained samples prepared as detailed above, in a Zeiss Elyra 7 Lattice SIM microscope, using a 63× 1.4 NA immersion oil objective. 3 colour Lattice SIM stacks were acquired with a 110 nm step size and reconstructed using the ZEN software (Zeiss). The final XY resolution of super-resolved images was 31.3 × 31.2 nm/px (2560 × 2560 px).

### Live imaging

Embryos were collected on apple juice agar plates with yeast paste, dechorionated in bleach and mounted in Voltalef medium (Samaro) between a semi-permeable membrane and a coverslip. The ventral side of the embryo was facing the coverslip.

Movies were acquired in a Leica SP8 confocal using a 40× apochromatic 1.3 objective, zoom ×2 and 400×400px size (providing an XY resolution of 0.36 μm/px), 12 bit depth, 400 Hz image acquisition frequency and pinhole of 4 airy units. In experiments where cellularization was quantified using the transmitted light channel, 33 × 2 μm slices were collected to reach the cross section of the embryo, providing a time resolution of 20 seconds per frame. In other experiments, 29 × 1 μm slices were collected, with total acquisition time of 15-60s per frame, depending on the experiment. Nup107-GFP movies were acquired using 4x zoom (0.18 μm/px in XY, 1 μm slices.)

### FRAP

Imaging of Notch-GFP and Dl-mScarlet during FRAP was performed as above, but using 4x zoom (0.18 μm/px XY resolution), 400×400 px size. Point bleaching was performed on 6 points targetting membranes per round of FRAP for 0.5s each (total bleach time 3 sec) simultaneously with 488 and 561nm laser. Pre and postbleaching images were collected at 400Hz (0.5 seconds / frame). FRAP was quantified by drawing circles of 20 px in diameter around the bleached regions and at another 6 control regions in non bleached membranes. FRAP recovery was calculated by dividing the average fluorescence at each region by the average pre-bleach intensity and normalized for the ratio of the average fluorescence at control regions to pre-bleach intensity, to account for loss of fluorescence due to bleaching during acquisition ***(Gomez-Lamarca et al. 2018)*.** Each curve was then scaled so that the first value after beaching was considered 0. Dl-mScarlet bleached too fast so FRAP recovery could not be accurately quantified.

### Image analysis

#### Quantifying membrane length

Length of membranes during cellularization was calculated from the orthogonal section in the center of the field of view. Fluorescent signal was thresholded using the Otsu method ***(Otsu 1979)*** and the height of the obtained object, equivalent to the length of the membrane at each time point, was calculated and plotted over time. Because the signal:noise ratio was different for each marker used, these quantifications were manually curated by marking the extent of membrane signal in the orthogonal view image when the automated segmentation did not match the raw signal.

#### Tracking nuclei and MS2 quantification

Movies were analyzed using custom MATLAB (MATLAB R2020a, MathWorks) scripts that have been previously described ***(Falo-Sanjuan et al. 2019)*,** with some adaptations.

Briefly, the His2Av-RFP signal was used to segment and track the nuclei in 3D. Each 3D stack was first filtered using a 3D median filter of 3 and increasing the contrast based on the intensity profile of each frame to account for bleaching. A Fourier transform log filter was then applied to enhance round objects ***(Garcia et al. 2013)*.** Segmentation was performed by applying a fixed intensity threshold to the filtered stack, which was empirically determined. This was followed by filters to fill holes in objects and discard miss-segmented nuclei based on size. 3D watershed accounting for anisotropic voxel sizes ***(Mishchenko 2015)*** was used to split merged nuclei. The final segmented stack was obtained by filtering by size again and and thickening each object. Lastly, the segmented stack was labelled to assign a number to each object, and the position of each centroid (in X, Y and Z) was calculated for tracking.

Nuclei were then tracked in 3D by finding the nearest object (minimum Euclidean distance between two centroids in space) in the previous 2 frames which was closer than 6 μm. If no object was found, that nucleus was kept with a new label. If more than one object was ‘matched’ with the same one in one of the previous 2 frames, both were kept with new labels.

After tracking, the positions of all pixels from each nucleus in each frame were used to measure the maximum fluorescence value in the GFP channel, which was used as a proxy of the spot fluorescence. Note that when a spot cannot be detected by eye this method detects only background, but the signal:background ratio is high enough that the subsequent analysis allows to classify confidently when the maximum value is really representing a spot.

#### Nuclear membrane tracking

To segment the nuclei in 3D from nuclear membrane markers (Nup107-GFP), each 3D stack was first resized to produce 1:1:1 ratio voxel sizes using the cubic interpolation from the *imresize3* function in MATLAB. Each resized stack was then filtered using a 3D gaussian filter of 1. To account for loss of fluorescence due to bleaching, the *imhistmatchn* function was used to adjust the contrast of each frame to the first one. A fixed intensity threshold of 10% was used to create a thresholded image, which was used as seed for Active Contour segmentation ***(Chan and Vese 2001)*** of the filtered image to produce an initial segmentation of nuclear membranes. The image was then inverted to recognize as object the space inside the nuclear membrane rather than the membrane itself. A filter based on the proportion of object present in each slice was used to remove the vitelline membrane. A 3D watershed filter was then used to separate merged objects and object thickening was used to compensate for any lost signal at the edges. Finally, 3D objects out of the range 10 μm^3^ to 200 μm^3^ were discarded. Segmented nuclei were then tracked in 3D as described in the previous section. In this case, because more nuclei were missing in each frame than when histones were segmented, a maximum distance of 4 μm was allowed for a nucleus to be considered the same as another in a maximum of 5 previous frames.

#### Nuclear 3D properties

After tracking, the MATLAB function *regionprops3* was used to extract 3D properties of each object: volume, surface area, solidity and length of principal axes. 2D slices at different fractions of the nuclear length (25, 50 and 75 %) were extracted and 2D properties quantified using *regionprops*: area, perimeter and eccentricity. Note the slices for 25, 50 and 75 % of the nuclear length were calculated at the level of the whole embryo, and due to the curvature, the measured properties will not correspond to the same position in all nuclei. Nevertheless, almost all the nuclei were imaged in the same plane so this is not expected to have a big influence in the 2D properties measured. The same approach to measure size and shape of nuclei was used using the histone rather than nucleoporin signal. In this case the fine details of the nuclear wrinkling could not be resolved but it provided a good approximation of the volume and length of the nuclei.

### Data processing and statistical analysis

#### MS2 data processing

Processing of MS2 data (definition of active nuclei and normalization for bleaching) has been carried out as described in our previous work ***(Falo-Sanjuan et al. 2019)*.** From the tracking step, the fluorescent trace of each nucleus over time was obtained. Only nuclei tracked for more than 10 frames were retained. First, nuclei were classified as inactive or active. To do so, the average of all nuclei (active and inactive) was calculated over time and fitted to a straight line. A median filter of 3 was applied to each nucleus over time to smooth the trace and ON periods were considered when fluorescent values were 1.2 times the baseline at each time point. This produced an initial segregation of active (nuclei ON for at least 5 frames) and inactive nuclei. These parameters were determined empirically on the basis that the filters retained nuclei with spots close to background levels and excluded false positives from bright background pixels. The mean fluorescence from MCP-GFP in the inactive nuclei was then used to define the background baseline and active nuclei were segregated again in the same manner. The final fluorescence values in the active nuclei were calculated by removing the fitted baseline from the maximum intensity value for each, and normalizing for the percentage that the MCP-GFP fluorescence in inactive nuclei decreased over time to account for the loss of fluorescence due to bleaching. Nuclei active in cycles before nc14 were discarded based on the timing of their activation.

In all movies, time into nc14 was considered from the end of the 13th syncytial division. Onsets of transcription were defined as the beginning of the first ON period, starting from 15 min into nc14 in most experiments, except for expression in the presence of maternal Gal4 (expression from 30 min to exclude earlier stochastic activity). The total mRNA output (in AU) was obtained by adding all the normalized transcription values for each cell in a defined time period. Cells producing ‘high’ and ‘low’ total mRNA output were defined by values that were above and below the median.

#### Statistical analysis

In figures and figure legends, n number indicates number of embryos imaged for each biological condition. Where appropriate, n number next to heatmaps indicates total number of cells combining all embryos for each biological condition. Plots showing mean levels of transcription and SEM (standard error of the mean) combine all traces from multiple embryos from the same biological condition.

### Reagents and software availability

Modifications in the existing code to track nuclei from the nuclear membrane signal and quantify nuclear morphology in 3D and 2D slices have been incorporated in a MATLAB app and can be obtained from https://github.com/BrayLab/LiveTrx.

## Movies

### Movie 1 - Changes in nuclear size and shape during nc14

Movie showing maximum projection of medial slices and orthogonal views of Nup107-GFP. 0.18 μm/px XY resolution and time resolution of 15s/frame. Anterior to the left; embryo imaged from the ventral side. Time indicates minutes from the beginning of nc14.

### Movie 2 - Localization of Notch and Delta during cellularization

Movie showing maximum projection of medial slices and orthogonal views of Dl-mScarlet (left) and Notch-GFP (right). 0.36 μm/px XY resolution and time resolution of 60s/frame. Anterior to the left; embryo imaged from the ventral side. Time indicates minutes from the beginning of nc14

### Movie 3 - Expression of *m5/m8* starts during cellularization

Movie showing cellularizing membranes using the marker Gap43-mCherry (maximum intensity projection of medial slices and orthogonal views, left) and transcription from *m5/m8-MS2 ^[II]^* (maximum intensity projection with maximum Y projection of the MCP-GFP channel, right). 0.36 μm/px XY resolution, 36×1mm slices and time resolution of 20s/frame. Anterior to the left; embryo imaged from the ventral side. Time indicates minutes from the beginning of nc14.

### Movie 4 - Expression of *m5/m8* in control, *slam* and *nullo* embryos

Movies showing MCP-GFP channel with transcription directed by *m5/m8 ^[III]^* (maximum intensity projection, left), His2Av-RFP channel in blue overlaid with MCP-GFP in green (maximum intensity projection, center) and transmitted light channel showing membrane growth (cross section, right) in control (top), *slam ^-/-^* (middle) and *nullo ^−^* (bottom) embryos. 0.36 μm/px XY resolution, 33×2mm slices and time resolution of 20s/frame. Anterior to the left; embryo imaged from the ventral side. Time indicates minutes from the beginning of nc14.

### Movie 5 - Expression of *m5/m8* in control and a*-Cat RNAi* embryos

Movies showing MCP-GFP channel with transcription directed by *m5/m8 ^[III]^* (maximum intensity projection, left) and His2Av-RFP channel in blue overlaid with MCP-GFP in green (maximum intensity projection, right) in control (top) and α-Cat depleted (bottom) embryos. 0.36 μm/px XY resolution, 32×1mm slices and time resolution of 20s/frame. Anterior to the left; embryo imaged from the ventral side. Time indicates minutes from the beginning of nc14.

### Movie 6 - Expression of *E(spl)m8-HLH* in control and α*-Cat RNAi* embryos

Movies showing MCP-GFP channel with *E(spl)m8-HLH* transcription (maximum intensity projection, left) and His2Av-RFP channel in blue overlaid with MCP-GFP in green (maximum intensity projection, right) in control (top) and α-Cat depleted (bottom) embryos. 0.36 μm/px XY resolution, 32×1mm slices and time resolution of 20s/frame. Anterior to the left; embryo imaged from the ventral side. Time indicates minutes from the beginning of nc14.

## Acknowledgments

We thank members of the Bray Lab for helpful discussions. Thanks to members of the Sanson lab for providing flies and advice and to Kat Millen and the Genetics Fly Facility for injections. We acknowledge the Cambridge Advanced Imaging Centre for their support, assistance in this work and use of their microscopes. This work was supported by a Wellcome Trust Investigator Award (212207/Z/18/Z) and a Medical Research Council Programme grant (MR/T014156/1) and by a PhD studentship to J.F.-S from the Wellcome Trust (109144/Z/15/Z).

## Author Contributions

J.F.-S. and S.J.B. planned the experiments; J.F.-S. conducted the experiments and analyzed the data; J.F.-S. and S.J.B. wrote the manuscript.

## Supplementary Information

**Figure S1.**
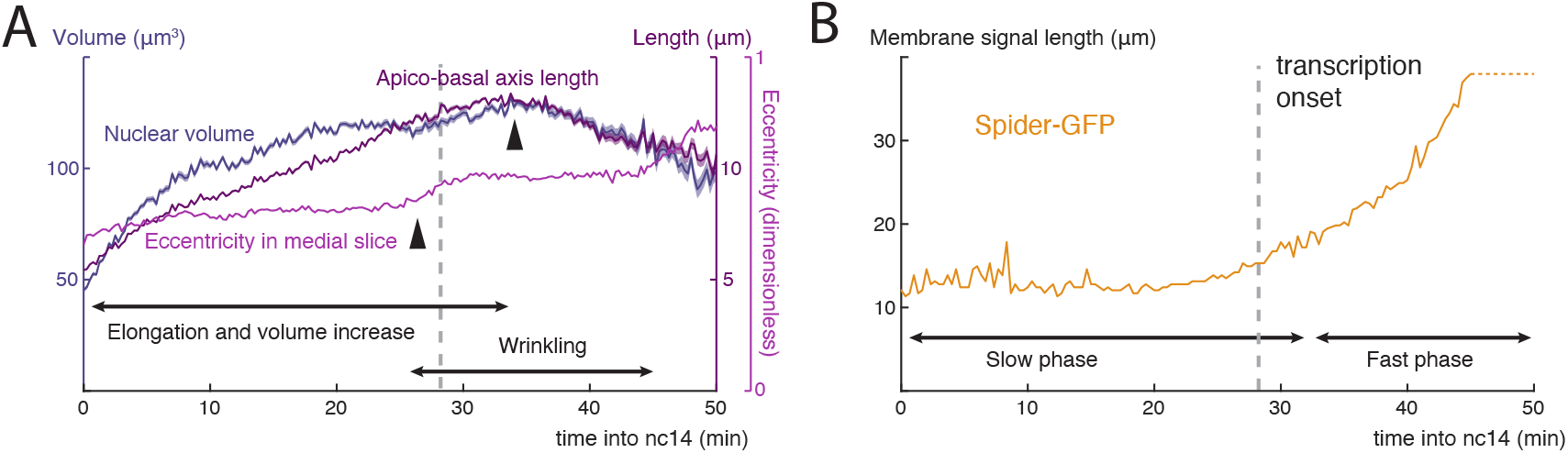
Correlation between developmental processes and onset of Notch dependent transcription. **A**) Timeline of changes in nuclear 3D properties over time, quantified using the nuclear membrane marker Nup107-GFP. **B**) Timeline of cellularization, measured by quantification of the length of the cell membrane marker Spider-GFP in orthogonal views. Dashed lines indicate length of signal is greater than the stack imaged.

**Figure S2.**
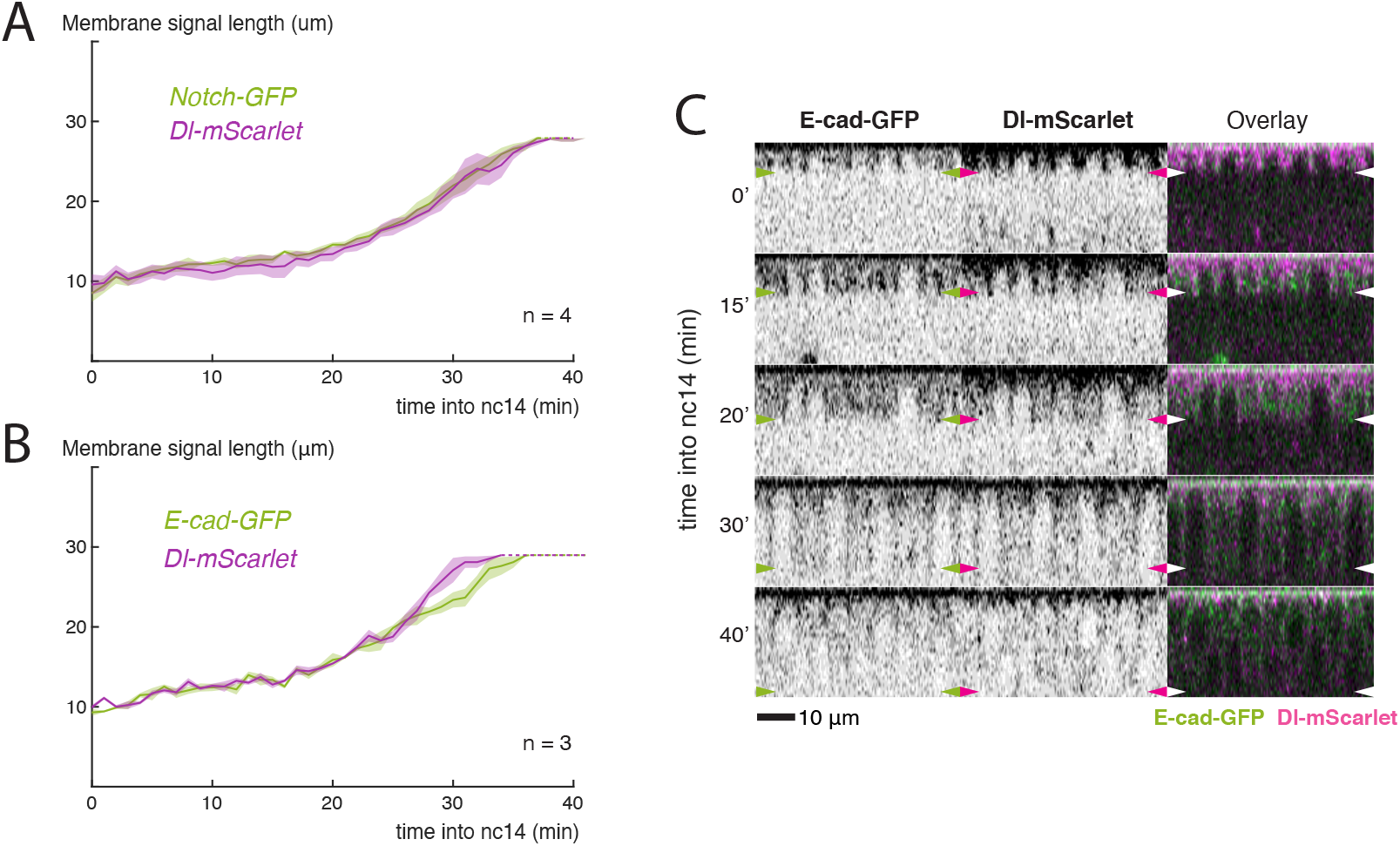
Delta tracks with E-cadherin as membranes grow. **A**) Comparison of the length of membrane occupied by Notch and Delta over time, extending basally at the same rate. **B**) Comparison of the length of lateral signal of E-cad and Delta over time. Dashed lines indicate membrane length is greater than the stack imaged. **C**) Orthogonal views from embryos expressing E-cad-GFP and Dl-mScarlet, showing colocalization at all timepoints during cellularization. Arrowheads indicate position of cellularization front.

**Figure S3.**
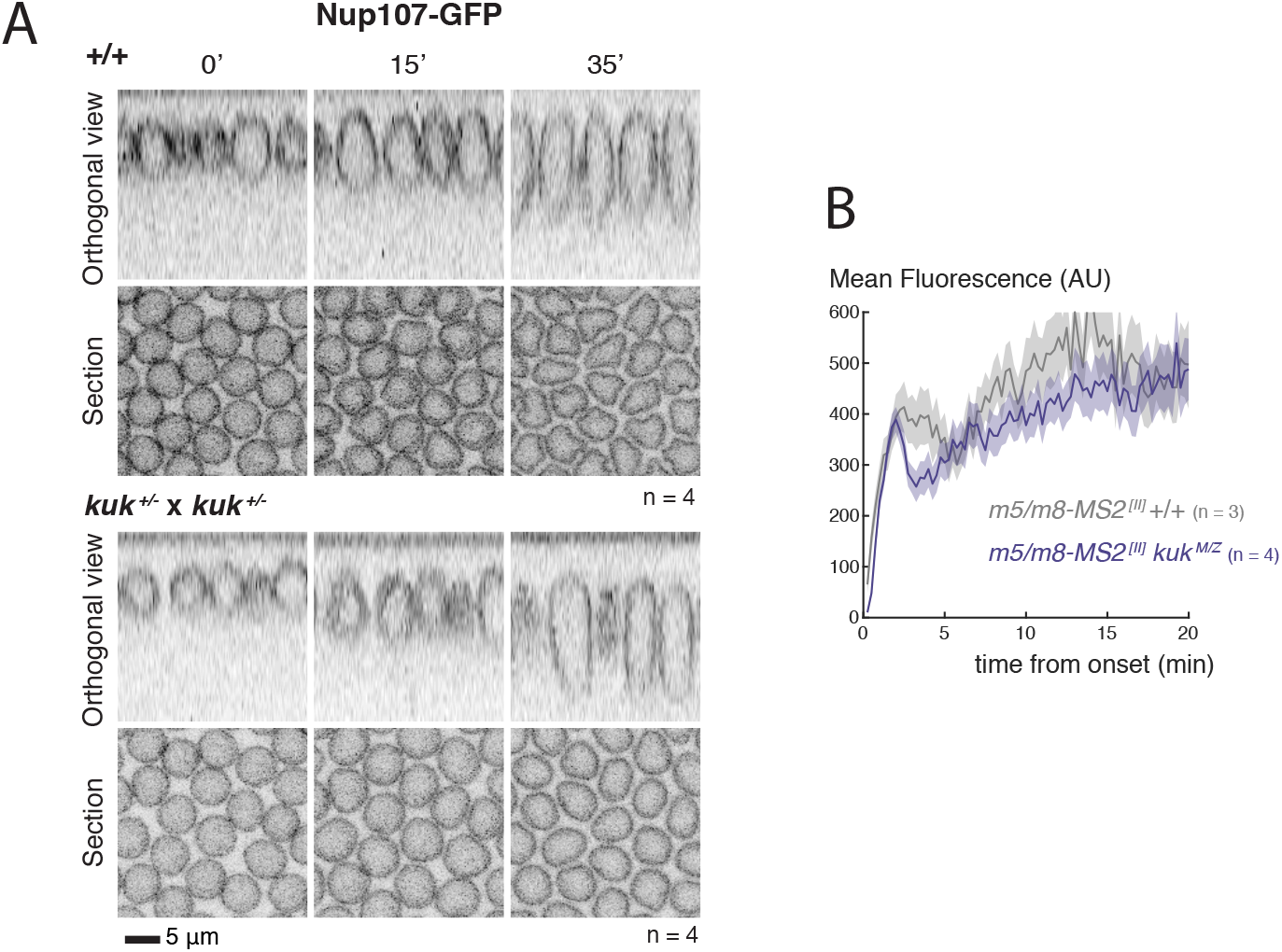
Changes in nuclear morphology do not influence Notch dependent transcription. **A**) Cross-sections and orthogonal views of the nuclear membrane marker Nup107-GFP in wild type (top) and embryos obtained from *kuk* heterozygous parents (bottom) at the indicated times (min into nc14), as this *kuk* allele was not homozygous viable in combination with Nup107-GFP. **B**) Mean levels of transcription when nuclei are aligned by onset times. Mean and SEM (shaded area) of all cells combined from multiple embryos are shown (n embryo numbers indicated in each).

**Figure S4.**
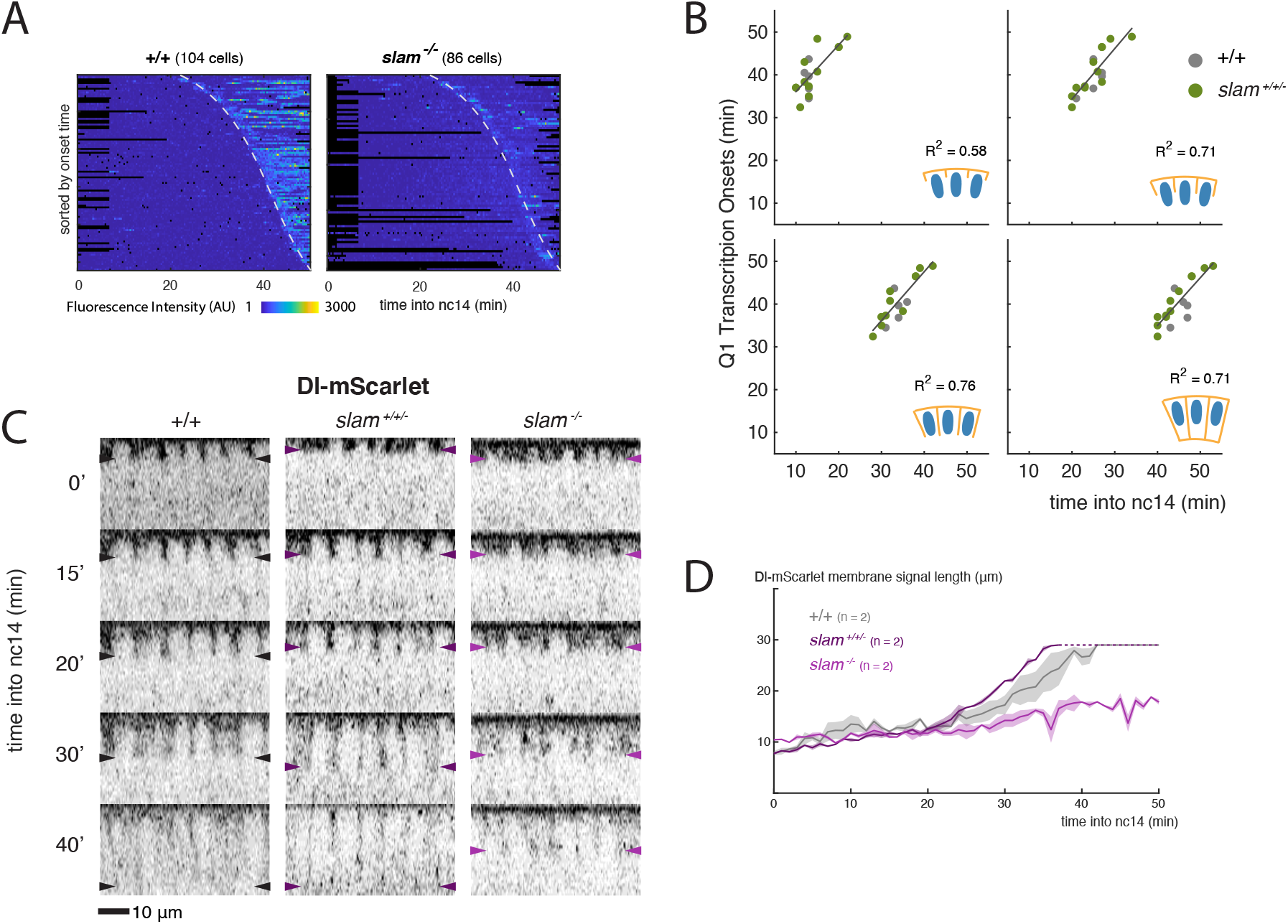
Delta localization in *slam* mutant embryos. **A**) Heatmaps of transcription in all mesectoderm nuclei from control and *slam^-/-^* embryos, sorted by onset time. Dashed lines indicate onset times in controls. **B**) Correlation between timepoints during cellularization (indicated by each cartoon) with onset of transcription from *m5/m8 ^[III]^* (calculated as the first quartile of onset times) in *slam^+/+/-^* and control embryos. R2 coefficients are calculated after pooling all points shown the same plot together. **C**) Orthogonal views from embryos expressing Dl-mScarlet in wild type, *slam^-/-^* or *slam^+/+/-^* backgrounds. Arrowheads indicate position of the most basal signal. **D**) Comparison of the length of membrane localization of Dl-mScarlet in wild type, *slam* homozygous embryos and other embryos obtained from the same cross (*slam^+/+/-^*). Delta did not extend basally in *slam^-/-^* embryos. Mean and SEM (shaded area), n embryos indicated for each. Dashed lines indicate membrane length is greater than the stack imaged.

**Figure S5.**
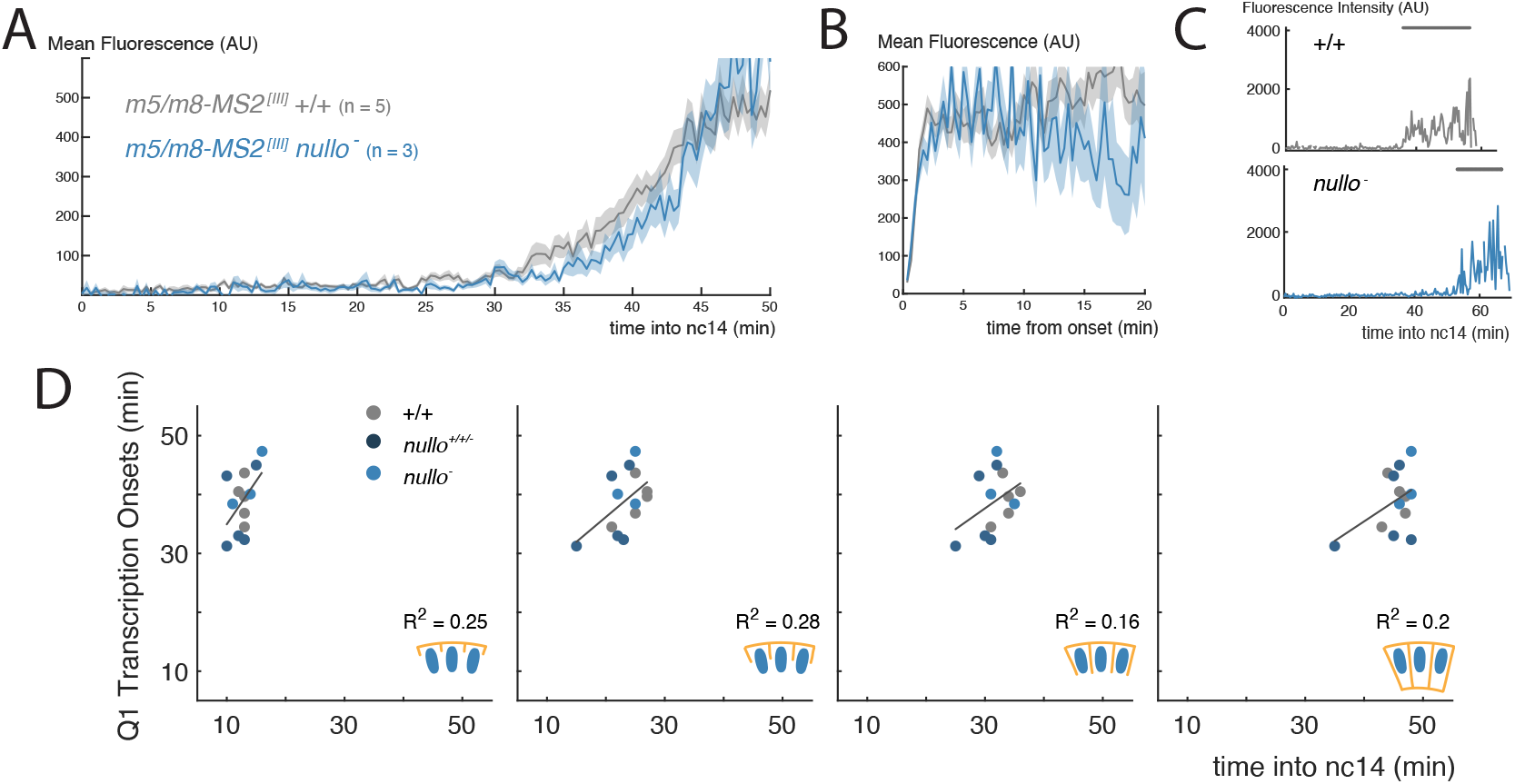
Absence of Nullo does not affect overall levels of transcription. **A**) Mean profile of *m5/m8 ^[III]^* activity in *nullo^−^* embryos compared to control embryos. **B**) Mean levels of transcription when nuclei are aligned by their onset times. **C**) Examples of transcription traces from mesectoderm nuclei. Grey lines indicate ON periods. **D**) Correlation between timepoints during cellularization (indicated by each cartoon) with onset of transcription from *m5/m8 ^[III]^* (calculated as the first quartile of onset times) in *nullo^+/+/-^*, *nullo^−^* and control embryos. R2 coefficients are calculated after pooling all points shown the same plot together. In **A** and **B**, mean and SEM (shaded area) of all cells combined from multiple embryos are shown (n embryo numbers indicated in each). Images, plots and quantifications of control embryos are duplicated from **Fig.** 4.

**Figure S6.**
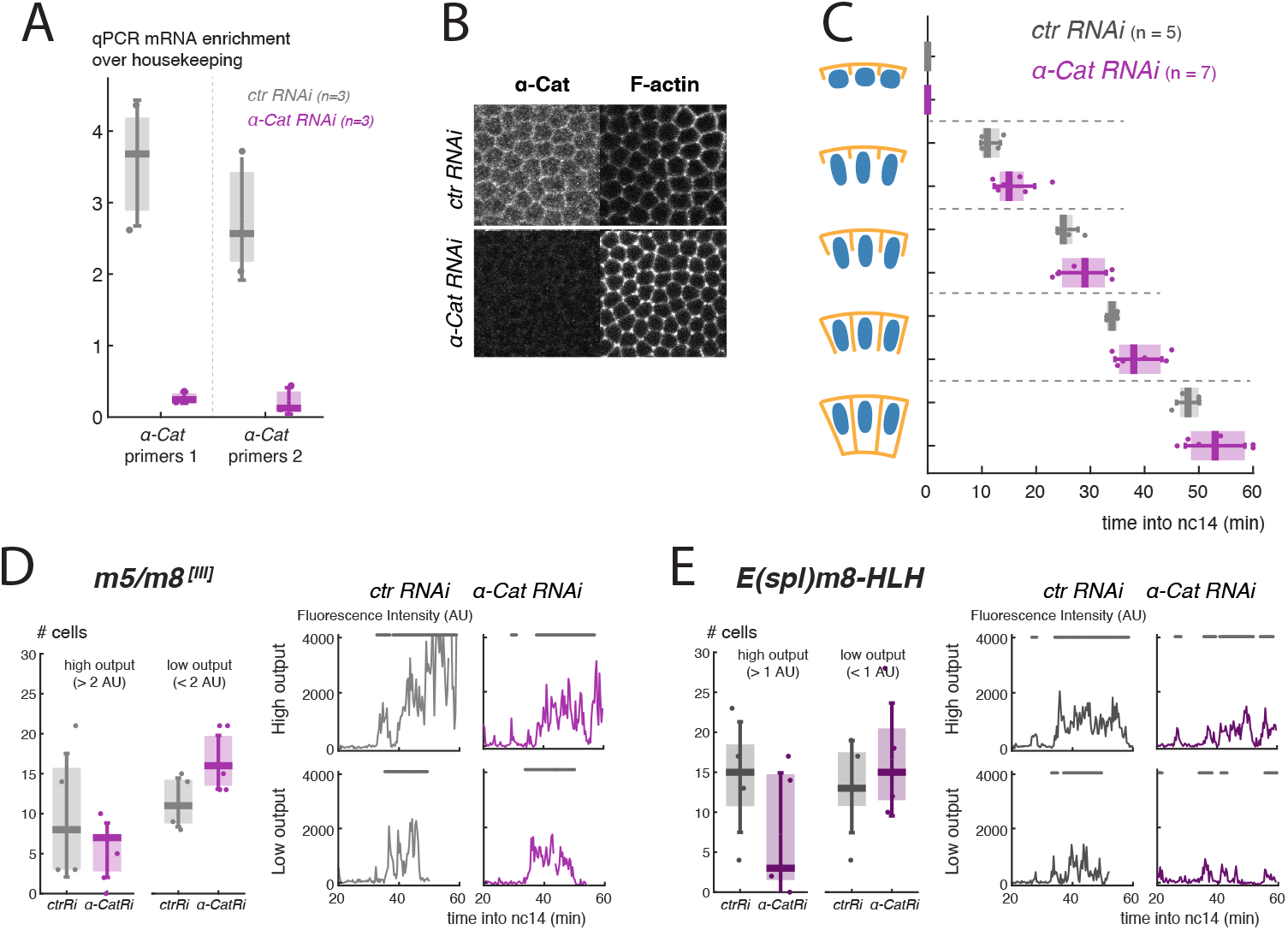
Adherens junctions influence Notch dependent transcription. **A**) Quantification of α*-Cat* mRNA levels by RT-qPCR (2 sets of primers) in pools of 15-20 eggs and/or pre-nc13 embryos upon control and α*-Cat* germline RNAi expression. n = 3 (control RNAi) and 3 (α*-Cat RNAi*) biological replicates. **B**) Mid-cellularization embryos stained for α-Cat and F-actin (phalloidin) upon control and α*-Cat* germline RNAi expression. **C**) Boxplots indicating timing of cellularization progression (timepoints when membranes reach each of the lengths with respect to nuclei indicated in the cartoons) in control and α*-Cat RNAi* conditions, quantified from *m5/m8 ^[III]^* MS2 movies. Median, Q1/Q3 quartiles and SD shown. **D**-**E**) Boxplots indicating number of cells producing high and low total levels of transcription (left, defined by production above and below the median) and examples of transcription traces from each group (right), for *m5/m8 ^[III]^* (**D**) and *E(spl)m8-HLH* (**E**). Grey lines indicate ON periods.

**Figure S7.**
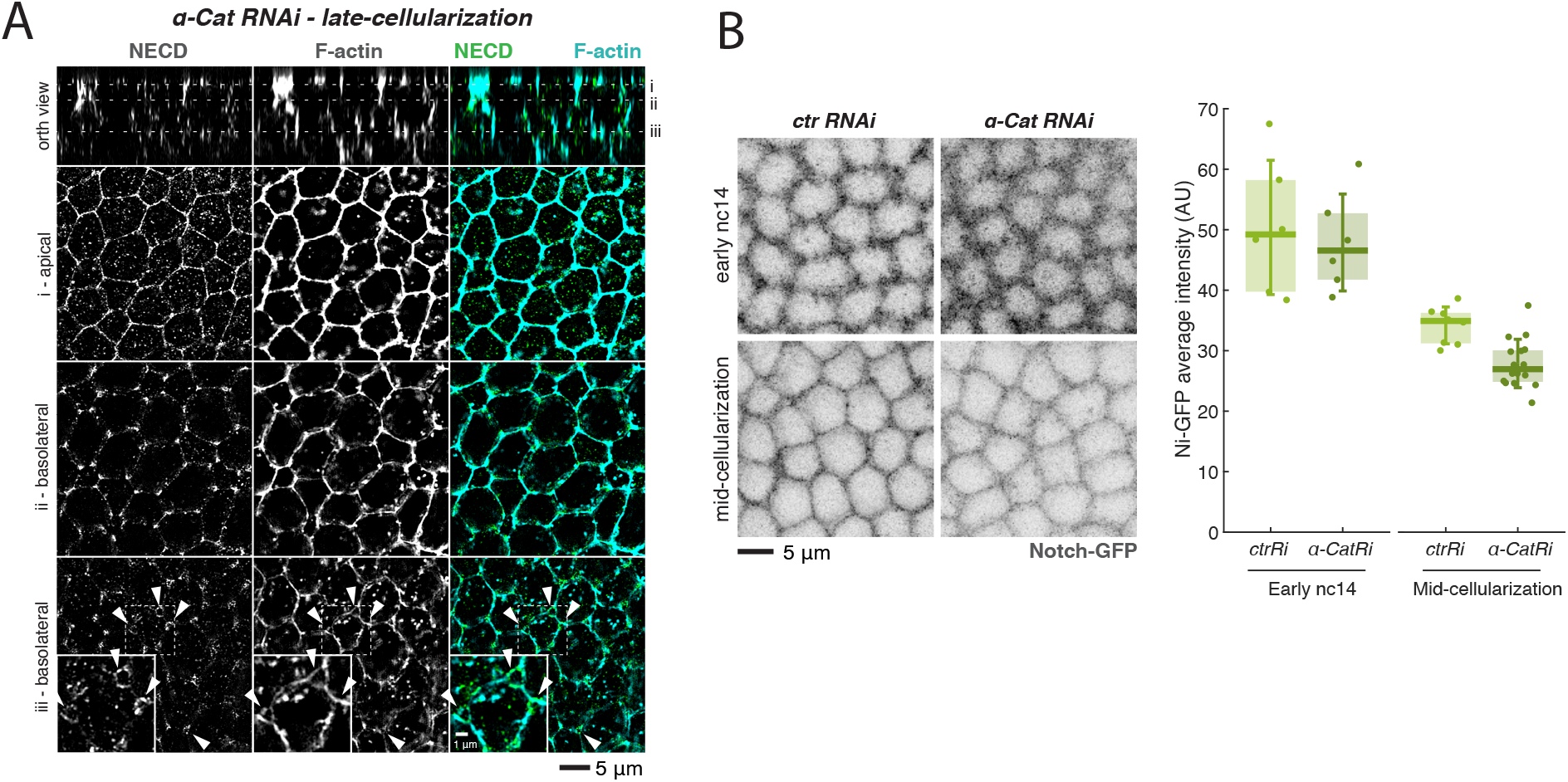
α-Catenin depletion does not influence Notch localization. **A**) Latecellularization α*-Cat* RNAi embryo stained with phalloidin and antibodies against NECD and E-cad and imaged using SIM (E-cad channel not shown). Arrowheads indicate holes in tricellular junctions caused by lack of adhesion. Top panels are orthogonal views with lines marking individual planes shown below. **B**) Stills of live early nc14 and mid-cellularization embryos expressing Notch-GFP upon control and α*-Cat* RNAi expression (left) and quantification of the overall Notch-GFP levels in each condition and timepoint (right). n = 6 (control RNAi early), 6 (α*-Cat* RNAi early), 8 (control RNAi mid-cellualarization) and 16 (α*-Cat* RNAi mid-cellularization).

## References

Bajpai, S., Correia, J., Feng, Y., Figueiredo, J., Sun, S.X., Longmore, G.D., Suriano, G., and Wirtz, D. (2008). α-Catenin mediates initial E-cadherin-dependent cell-cell recognition and subsequent bond strengthening. Proc. Natl. Acad. Sci. 105.47, 18331–18336.

Bothma, J.P., Garcia, H.G., Esposito, E., Schlissel, G., Gregor, T., and Levine, M. (2014). Dynamic regulation of eve stripe 2 expression reveals transcriptional bursts in living Drosophila embryos. Proc. Natl. Acad. Sci. 111.29, 10598–10603.

Bothma, J.P., Garcia, H.G., Ng, S., Perry, M.W., Gregor, T., and Levine, M. (2015). Enhancer additivity and non-additivity are determined by enhancer strength in the Drosophila embryo. Elife 4.AUGUST2015, 1–14.

Boukhatmi, H., Martins, T., Pillidge, Z., Kamenova, T., and Bray, S. (2020). Notch Mediates Inter-tissue Communication to Promote Tumorigenesis. Curr. Biol. 30.10, 1809–1820.e4.

Brandt, A., Papagiannouli, F., Wagner, N., Wilsch-Bräuninger, M., Braun, M., Furlong, E.E., Loserth, S., Wenzl, C., Pilot, F., Vogt, N., Lecuit, T., Krohne, G., and Großhans, J. (2006). Developmental Control of Nuclear Size and Shape by kugelkern and kurzkern. Curr. Biol. 16.6, 543–552.

Chan, T. and Vese, L. (2001). Active contours without edges. IEEE Trans. Image Process. 10.2, 266–277.

Coumailleau, F., Fürthauer, M., Knoblich, J.A., and González-Gaitán, M. (2009). Directional Delta and Notch trafficking in Sara endosomes during asymmetric cell division. Nature 458.7241, 1051–1055.

Couturier, L., Vodovar, N., and Schweisguth, F. (2012). Endocytosis by Numb breaks Notch symmetry at cytokinesis. Nat. Cell Biol. 14.2, 131–139.

Cowden, J. and Levine, M. (2002). The Snail repressor positions Notch signaling in the Drosophila embryo. Development 129.7, 1785–93.

De Renzis, S., Yu, J., Zinzen, R., and Wieschaus, E. (2006). Dorsal-ventral pattern of Delta trafficking is established by a snail-tom-neuralized pathway. Dev. Cell 10.2, 257–264.

Fabrowski, P., Necakov, A.S., Mumbauer, S., Loeser, E., Reversi, A., Streichan, S., Briggs, J.A.G., and De Renzis, S. (2013). Tubular endocytosis drives remodelling of the apical surface during epithelial morphogenesis in Drosophila. Nat. Commun. 4.1, 2244.

Falo-Sanjuan, J., Lammers, N.C., Garcia, H.G., and Bray, S.J. (2019). Enhancer Priming Enables Fast and Sustained Transcriptional Responses to Notch Signaling. Dev. Cell 50.4, 411–425.e8.

Foe, V.E. and Alberts, B.M. (1983). Studies of nuclear and cytoplasmic behaviour during the five mitotic cycles that precede gastrulation in Drosophila embryogenesis. J. Cell Sci. 61, 31–70.

Garcia, H.G., Tikhonov, M., Lin, A., and Gregor, T. (2013). Quantitative Imaging of Transcription in Living Drosophila Embryos Links Polymerase Activity to Patterning. Curr. Biol. 23.21, 2140–2145.

Gomez-Lamarca, M.J., Falo-Sanjuan, J., Stojnic, R., Abdul Rehman, S., Muresan, L., Jones, M.L., Pillidge, Z., Cerda-Moya, G., Yuan, Z., Baloul, S., Valenti, P., Bystricky, K., Payre, F., O’Holleran, K., Kovall, R., and Bray, S.J. (2018). Activation of the Notch Signaling Pathway In Vivo Elicits Changes in CSL Nuclear Dynamics. Dev. Cell 44.5, 611–623.e7.

Gordon, W.R., Zimmerman, B., He, L., Miles, L.J., Huang, J., Tiyanont, K., McArthur, D.G., Aster, J.C., Perrimon, N., Loparo, J.J., and Blacklow, S.C. (2015). Mechanical Allostery: Evidence for a Force Requirement in the Proteolytic Activation of Notch. Dev. Cell 33.6, 729–736.

Hampoelz, B., Mackmull, M.-T., Machado, P., Ronchi, P., Bui, K.H., Schieber, N., Santarella-Mellwig, R., Necakov, A., Andrés-Pons, A., Philippe, J.M., Lecuit, T., Schwab, Y., and Beck, M. (2016). Pre-assembled Nuclear Pores Insert into the Nuclear Envelope during Early Development. Cell 166.3, 664–678.

Hatakeyama, J., Wakamatsu, Y., Nagafuchi, A., Kageyama, R., Shigemoto, R., and Shimamura, K. (2014). Cadherin-based adhesions in the apical endfoot are required for active Notch signaling to control neurogenesis in vertebrates. Development 141.8, 1671–1682.

Hong, J.W., Park, K.W., and Levine, M.S. (2013). Temporal regulation of single-minded target genes in the ventral midline of the Drosophila central nervous system. Dev. Biol. 380.2, 335–343.

Hoppe, C., Bowles, J.R., Minchington, T.G., Sutcliffe, C., Upadhyai, P., Rattray, M., and Ashe, H.L. (2020). Modulation of the Promoter Activation Rate Dictates the Transcriptional Response to Graded BMP Signaling Levels in the Drosophila Embryo. Dev. Cell 54.6, 727–741.e7.

Huang, H. and Kornberg, T.B. (2015). Myoblast cytonemes mediate Wg signaling from the wing imaginal disc and Delta-Notch signaling to the air sac primordium. Elife 4.MAY, 1–22.

Huang, J., Zhou, W., Dong, W., Watson, A.M., and Hong, Y. (2009). Directed, efficient, and versatile modifications of the Drosophila genome by genomic engineering. Proc. Natl. Acad. Sci. 106.20, 8284–8289.

Hunter, C., Sung, P., Schejter, E.D., and Wieschaus, E. (2002). Conserved Domains of the Nullo Protein Required for Cell-Surface Localization and Formation of Adherens Junctions. Mol. Biol. Cell 13.1, 146–157.

Hunter, C. and Wieschaus, E. (2000). Regulated Expression of nullo Is Required for the Formation of Distinct Apical and Basal Adherens Junctions in the Drosophila Blastoderm. J. Cell Biol. 150.2, 391–402.

Hunter, G.L., He, L., Perrimon, N., Charras, G., Giniger, E., and Baum, B. (2019). A role for actomyosin contractility in Notch signaling. BMC Biol. 17.1, 1–15.

Ishiyama, N., Sarpal, R., Wood, M.N., Barrick, S.K., Nishikawa, T., Hayashi, H., Kobb, A.B., Flozak, A.S., Yemelyanov, A., Fernandez-Gonzalez, R., Yonemura, S., Leckband, D.E., Gottardi, C.J., Tepass, U., and Ikura, M. (2018). Force-dependent allostery of the α-catenin actin-binding domain controls adherens junction dynamics and functions. Nat. Commun. 9.1, 1–17.

Izquierdo, E., Quinkler, T., and De Renzis, S. (2018). Guided morphogenesis through optogenetic activation of Rho signalling during early Drosophila embryogenesis. Nat. Commun. 9.1, 2366.

Jurado, J., de Navascués, J., and Gorfinkiel, N. (2016). α-Catenin stabilises Cadherin-Catenin complexes and modulates actomyosin dynamics to allow pulsatile apical contraction. J. Cell Sci. 129.24, 4496–4508.

Katsani, K.R., Karess, R.E., Dostatni, N., and Doye, V. (2008). In Vivo Dynamics of Drosophila Nuclear Envelope Components. Mol. Biol. Cell 19.1, 3652–3666.

Khait, I., Orsher, Y., Golan, O., Binshtok, U., Gordon-Bar, N., Amir-Zilberstein, L., and Sprinzak, D. (2016). Quantitative Analysis of Delta-like 1 Membrane Dynamics Elucidates the Role of Contact Geometry on Notch Signaling. Cell Rep. 14.2, 225–233.

Kramer, H. (2000). The ups and downs of life in an epithelium. J. Cell Biol. 151.4, 15–18.

Kwak, M., Southard, K., Kim, N.H., Gopalappa, R., Kim, W.R., An, M., Lee, H.J., Farlow, J., Georgakopoulos, A., Robakis, N., Seo, D., Kim, H.B., Kim, Y.H., Cheon, J., Gartner, Z., and Jun, Y.-w. (2020). Size-dependent protein segregation creates a spatial switch for Notch and APP signaling. bioRxiv, 1–25.

Lecuit, T., Samanta, R., and Wieschaus, E. (2002). slam Encodes a Developmental Regulator of Polarized Membrane Growth during Cleavage of the Drosophila Embryo. Dev. Cell 2.4, 425–436.

Lecuit, T. and Wieschaus, E. (2000). Polarized Insertion of New Membrane from a Cytoplasmic Reservoir during Cleavage of the Drosophila Embryo. J. Cell Biol. 150.4, 849–860.

Lim, B., Levine, M., and Yamazaki, Y. (2017). Transcriptional Pre-patterning of Drosophila Gastrulation. Curr. Biol. 27.2, 286–290.

López-Schier, H. and St Johnston, D. (2001). Delta signaling from the germ line controls the proliferation and differentiation of the somatic follicle cells during Drosophila oogenesis. Genes Dev. 15.11, 1393–405.

MartÍn-Bermudo, M.D., Carmena, A., and Jiménez, F. (1995). Neurogenic genes control gene expression at the transcriptional level in early neurogenesis and in mesectoderm specification. Development 121.1, 219–224.

Mishchenko, Y. (2015). A fast algorithm for computation of discrete Euclidean distance transform in three or more dimensions on vector processing architectures. Signal, Image Video Process. 9.1, 19–27.

Morel, V., Le Borgne, R., and Schweisguth, F. (2003). Snail is required for Delta endocytosis and Notch-dependent activation of single-minded expression. Dev. Genes Evol. 213.2, 65–72.

Morel, V. and Schweisguth, F. (2000). Repression by Suppressor of Hairless and activation by Notch are required to define a single row of single-minded expressing cells in the Drosophila embryo. Genes Dev. 14.3, 377–388.

Morin, X., Daneman, R., Zavortink, M., and Chia, W. (2001). A protein trap strategy to detect GFP-tagged proteins expressed from their endogenous loci in Drosophila. Proc. Natl. Acad. Sci. U. S. A. 98.26, 15050–15055.

Nambu, J.R., Franks, R.G., Hu, S., and Crews, S.T. (1990). The single-minded gene of Drosophila is required for the expression of genes important for the development of CNS midline cells. Cell 63.1, 63–75.

Nandagopal, N., Santat, L.A., and Elowitz, M.B. (2019). Cis-activation in the Notch signaling pathway. Elife 8, 1–34.

Otsu, N. (1979). A Threshold Selection Method from Gray-Level Histograms. IEEE Trans. Syst. Man. Cybern. 9.1, 62–66.

Pilot, F., Philippe, J.-M., Lemmers, C., Chauvin, J.-P., and Lecuit, T. (2006). Developmental control of nuclear morphogenesis and anchoring by charleston, identified in a functional genomic screen of Drosophila cellularisation. Development 133.4, 711–23.

Shaya, O., Binshtok, U., Hersch, M., Rivkin, D., Weinreb, S., Amir-Zilberstein, L., Khamaisi, B., Oppenheim, O., Desai, R.A., Goodyear, R.J., Richardson, G.P., Chen, C.S., and Sprinzak, D. (2017). Cell-Cell Contact Area Affects Notch Signaling and Notch-Dependent Patterning. Dev. Cell 40.5, 505–511.e6.

Sokac, A.M. and Wieschaus, E. (2008a). Zygotically controlled F-actin establishes cortical compartments to stabilize furrows during Drosophila cellularization. J. Cell Sci. 121.11, 1815–1824.

Sokac, A.M. and Wieschaus, E. (2008b). Local Actin-Dependent Endocytosis Is Zygotically Controlled to Initiate Drosophila Cellularization. Dev. Cell 14.5, 775–786.

Staller, M.V., Yan, D., Randklev, S., Bragdon, M.D., Wunderlich, Z.B., Tao, R., Perkins, L.A., DePace, A.H., and Perrimon, N. (2013). Depleting Gene Activities in Early Drosophila Embryos with the “Maternal-Gal4–shRNA” System. Genetics 193.1, 51–61.

Trylinski, M., Mazouni, K., and Schweisguth, F. (2017). Intra-lineage Fate Decisions Involve Activation of Notch Receptors Basal to the Midbody in Drosophila Sensory Organ Precursor Cells. Curr. Biol. 27.15, 2239–2247.e3.

Viswanathan, R., Necakov, A., Trylinski, M., Harish, R.K., Krueger, D., Esposito, E., Schweisguth, F., Neveu, P., and De Renzis, S. (2019). Optogenetic inhibition of Delta reveals digital Notch signalling output during tissue differentiation. EMBO Rep. 20.12, e47999.

Wenzl, C., Yan, S., Laupsien, P., and Großhans, J. (2010). Localization of RhoGEF2 during Drosophila cellularization is developmentally controlled by slam. Mech. Dev. 127.7–8, 371–384.

Yu, H.H. and Zallen, J.A. (2020). Abl and Canoe/Afadin mediate mechanotransduction at tricellular junctions. Science (80-.). 370.6520, eaba5528.

Zinzen, R.P., Cande, J., Ronshaugen, M., Papatsenko, D., and Levine, M. (2006). Evolution of the Ventral Midline in Insect Embryos. Dev. Cell 11.6, 895–902.

